# Opposing Wnt and JAK-STAT signaling gradients define a stem cell domain by regulating spatially patterned cell division and differentiation at two borders

**DOI:** 10.1101/2020.06.23.167536

**Authors:** David Melamed, Daniel Kalderon

**Affiliations:** Dept. of Biological Sciences, Columbia University, New York

## Abstract

Many adult stem cells are maintained as a community by population asymmetry, wherein stochastic actions of individual cells collectively result in a balance between stem cell division and differentiation. We have used Drosophila Follicle Stem Cells (FSCs) as a paradigm to explore the extracellular niche signals that define a stem cell domain and organize stem cell behavior. FSCs produce transit-amplifying Follicle Cells (FCs) from their posterior face and quiescent Escort Cells (ECs) to their anterior. Here we show that JAK-STAT pathway activity, which declines from posterior to anterior, dictates the pattern of divisions over the FSC and EC domains, promotes more posterior FSC locations and conversion to FCs, while opposing EC production. A Wnt pathway gradient of opposite polarity promotes more anterior FSC locations and EC production and opposes FC production. Promotion of both FSC division and conversion to FCs by JAK-STAT signaling buffers the effects of genetically altered pathway activity on FSC numbers and balances the four-fold higher rate of differentiation at the posterior face of the FSC domain with a higher rate of FSC division in the most posterior layer. However, genetic elimination of Wnt pathway activity exacerbated elevated FC production resulting from increased JAK-STAT pathway activity, leading to rapid FSC depletion despite high rates of division. The two pathways combine to define a stem cell domain through concerted effects on FSC differentiation to ECs (high Wnt, low JAK-STAT) and FCs (low Wnt, high JAK-STAT) at each end of opposing signaling gradients, further enforced by quiescence at the anterior border due to declining JAK-STAT pathway activity.

## Introduction

The physiological role of adult stem cells is to maintain appropriate production of a restricted set of cell types throughout life (Clevers and Watt, 2018; Post and Clevers, 2019). To accomplish this objective, a sufficient population of stem cells must itself be maintained. Consequently, there must be some mechanism that balances stem cell proliferation and differentiation. The balance need not be precise or without fluctuations, especially if the stem cell population is large and not at risk of temporary or long-term extinction. If the number of stem cells is held roughly constant over time, then an unchanging anatomy can provide a constant environment for regulating stem cell divisions and guiding differentiation of their products.

The balance between stem cell division and differentiation can operate at the single cell level or at the community level (Jones, 2010; Mesa et al., 2018; Snippert et al., 2010). If each stem cell repeatedly divides to produce a stem cell and a differentiated product (“single cell asymmetry”), the rate of division must simply be matched to the required supply of product cells. More commonly, however, a group of stem cells in a given location is maintained by “population asymmetry”, where individual stem cells exhibit non-uniform, stochastic behaviors and differentiation need not be mechanistically linked to division of the same stem cell (Reilein et al., 2018; Ritsma et al., 2014; Rompolas et al., 2016; Simons and Clevers, 2011). The behavior of such stem cells is substantially guided by extracellular signals, raising the important questions of how such signals might define niche space and the number of stem cells accommodated, how they affect stem cell division and differentiation, and whether they co-ordinate those two fundamental behaviors. Drosophila ovarian Follicle Stem Cells (FSCs) provide an outstanding paradigm to pursue these questions.

FSCs were first defined as the source cells for the Follicle Cell (FC) epithelium that surrounds each egg chamber. An egg chamber buds every 12h under optimal conditions, from the germarium of each of a female’s thirty or more ovarioles (Fig. 1A-D), requiring a high constitutive rate of FC production (Duhart et al., 2017; Margolis and Spradling, 1995). Production of 5-6 “founder” FCs (proliferative precursors of mature FCs) per budding cycle is accomplished by 14-16 FSCs, arranged in three anterior-posterior (AP) rings near the mid-point of the germarium (Fig. 1A) (Hayashi et al., 2020; Reilein et al., 2017; Reilein et al., 2018). These FSCs also produce a second cell type known as an Escort Cell (EC) (Hayashi et al., 2020; Reilein et al., 2017). ECs are quiescent cells anterior to the FSC domain (Fig. 1A) that envelop and support the differentiation of developing germline cysts (Decotto and Spradling, 2005; Kirilly et al., 2011). FCs, which first encapsulate region 2b germline cysts and are defined by continued association with a single cyst, derive directly from the posterior (“layer 1”) FSCs, whereas ECs derive directly from anterior FSCs (Reilein et al., 2017). Each FSC lineage (marked descendants of a single FSC) can undergo rapid extinction or amplification in an apparently stochastic manner, and each lineage can include both ECs and FCs because FSCs can exchange AP locations; they are also radially mobile (Reilein et al., 2017). Posterior FSCs divide faster than anterior FSCs, so that the greater efflux from the posterior face of the FSC domain is supported without significant net flow of FSCs from anterior to posterior locations (Reilein et al., 2017). FSC division and differentiation to a FC are separate events; they are not temporally or functionally coupled in an individual FSC (Reilein et al., 2018). Thus, FSCs present a paradigm of dynamic heterogeneity (Greulich and Simons, 2016), wherein a community of stem cells exists within a defined spatial domain and constituent stem cells exhibit both distinctive instantaneous properties according to precise AP location and fluid, apparently stochastic changes in position and behavior over time. How are these heterogeneous behaviors marshaled into a defined stem cell domain that maintains a constant number of stem cells and a continuous supply of an appropriate number of FC and EC products?

**Figure 1.**
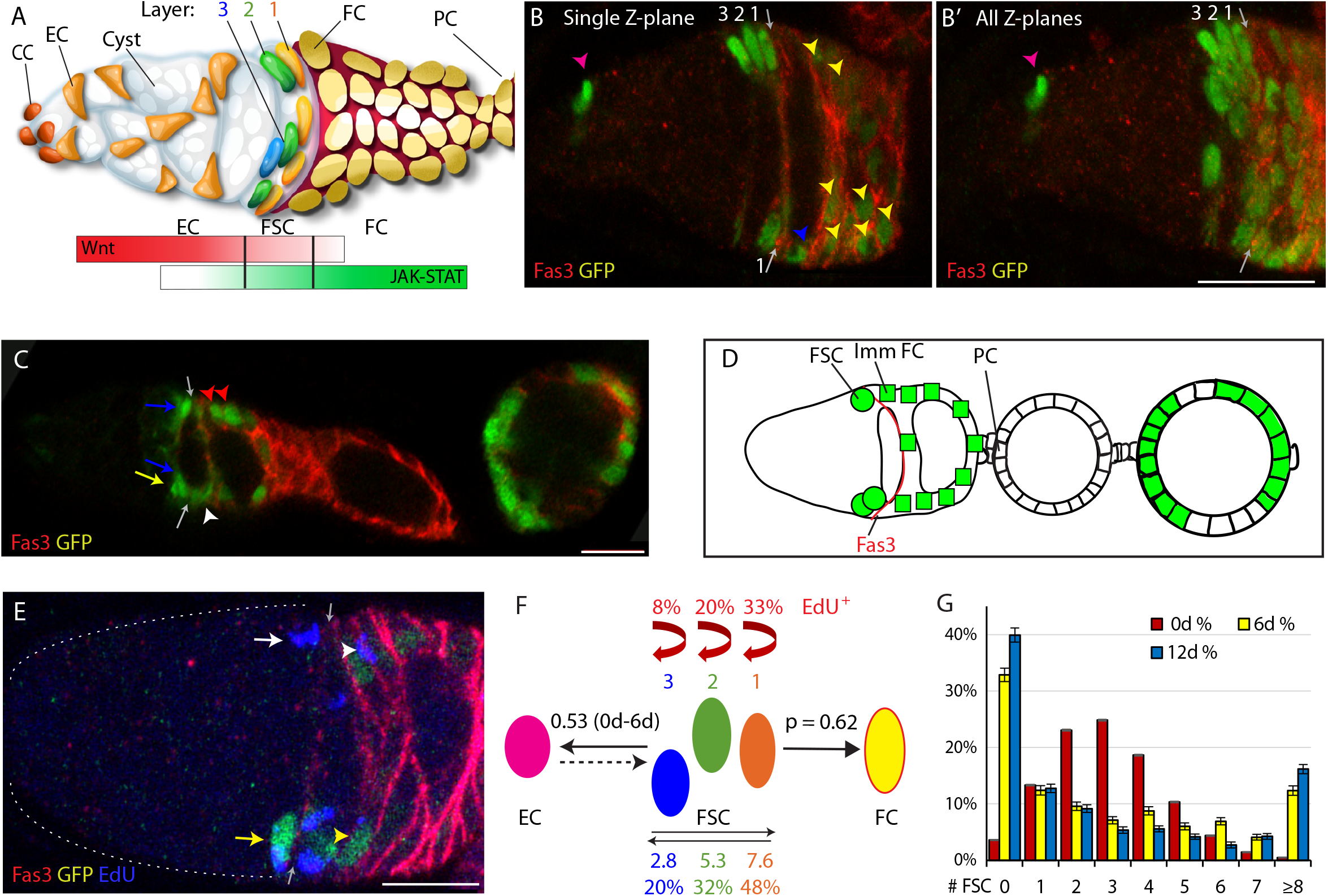
Follicle Stem Cell locations, signals and behaviors. (A) Cartoon representation of a germarium. Cap Cells (CC) at the anterior (left) contact Germline Stem Cells (not shown), which produce Cystoblast daughters that mature into 16-cell germline cysts (white) as they progress posteriorly. Quiescent Escort Cells (EC) extend processes around germline cysts and support their differentiation. Follicle Stem Cells (FSCs) occupy three AP Layers (3, 2, 1) around the germarial circumference and immediately anterior to strong Fas3 staining (red) on the surface of all early Follicle Cells (FC). FCs proliferate to form a monolayer epithelium, including specialized terminal Polar Cells (PC), which secrete the Upd ligand responsible for generating a JAK-STAT pathway gradient (green) of opposite polarity to the Wnt pathway gradient (red), generated by ligands produced in CCs and ECs. (B) A GFP-positive (green) MARCM FSC lineage that includes FSCs in each layer, an EC (magenta arrowhead), a recently produced “immediate” FC (blue arrowhead) and other FCs (yellow arrowheads) visualized together with Fas3 (red, arrows mark anterior border) as (B) a single 3μm z-section and (B’) a projection of ten z-sections (scoring is done by examining each z-section). (C, D) Early portion of an ovariole with marked FSCs (green) in layer 1 (blue arrows) and layer 2 (yellow arrows), a marked immediate FC (blue arrowhead) and more posterior FCs (yellow arrowheads) together with the anterior Fas3 (red) border (gray arrows), also (D) shown diagrammatically with anterior Polar Cells (PC) of the first budded egg chamber indicated. (E) Germarium with a MARCM FSC lineage (green) stained for EdU (blue) incorporation during 1h prior to fixation, showing examples of a GFP-positive EdU^+^ FSC (yellow arrow) and FC (yellow arrowhead), a GFP-negative EdU^+^ FSC (white arrow) and FC (white arrowhead) and the anterior Fas3 (red) border (gray arrows). (F) Diagram showing four of five properties of FSC behavior measured for all marked FSCs in MARCM lineages, with values indicated for normal FSCs: EdU incorporation frequency for each FSC layer (red text and arrows), FSC location among the three layers (indicated by absolute numbers and frequencies), ECs produced per anterior FSC over a given period (0.53 from 0-6d), and the likelihood for a layer 1 FSC to become an FC (p = 0.62) in a single budding cycle. (G) Distribution of the number of surviving FSCs observed for control genotypes at 6d (yellow) and 12d (blue), with the theoretically expected binomial distribution of FSCs initially marked (0d; red) based on the measured average number of surviving marked FSCs. All scale bars are 10μm.

FSC maintenance and amplification have been found to depend on the activity of many of the major pathways initiated by extracellular signals. The earliest studies highlighted the role of Hedgehog (Hh) signaling (Zhang and Kalderon, 2001). Hh was shown to influence FSC behavior principally by regulating the rate of FSC division through transcriptional induction of the Hippo-pathway transcriptional co-activator Yorkie (Yki) (Huang and Kalderon, 2014). The key role of FSC division rate for FSC competition was highlighted also by the discovery of several regulators of proliferation in a genetic screen for FSC maintenance factors (Wang et al., 2012; Wang and Kalderon, 2009), and was explained by the finding that FSC differentiation is independent of FSC division (i.e., these are two separate processes) (Reilein et al., 2018). Although the Hh signal is graded, declining from anterior to posterior, initial tests indicated that graded signaling was not important for continued normal FSC function (Vied et al., 2012). BMP, EGF, integrin and Insulin Receptor initiated pathways have also been implicated in FSC function (Castanieto et al., 2014; Johnston et al., 2016; Kirilly et al., 2005; O’Reilly et al., 2008; Vied et al., 2012; Wang et al., 2012) but the two pathways that have emerged so far as likely determinants of niche space and position-specific stem cell behaviors are the Wnt and JAK-STAT pathways because they both have graded activities in the AP dimension (Fig. 1A) (Reilein et al., 2017; Vied et al., 2012; Wang and Page-McCaw, 2014) and both have a very strong influence on FSC behavior (Reilein et al., 2017; Song and Xie, 2003; Vied et al., 2012).

The *wg* and Wnt6 ligands are produced in Cap Cells at the anterior of the germarium and are supplemented by the production of Wnt2 and Wnt4 in ECs (Forbes et al., 1996; Luo et al., 2015; Sahai-Hernandez and Nystul, 2013; Waghmare and Page-McCaw, 2018; Wang and Page-McCaw, 2018) to produce high levels of pathway activity over the EC domain with a sharp decline over the FSC domain and no detectable activity in FCs (Fig. 1A) (Reilein et al., 2017; Wang and Page-McCaw, 2014). FSCs were lost from the niche cell autonomously when Wnt signaling was either genetically elevated or reduced (Song and Xie, 2003; Vied et al., 2012). More recently it was shown that the primary effects of altering Wnt pathway activity were exerted on the AP location of FSCs and their conversion to differentiated products, with increased pathway activity favoring more anterior locations and EC production, while reducing FC production (Reilein et al., 2017). Thus, relatively rapid loss of FSCs due to elevated Wnt pathway activity results from conversion of all FSCs over time to ECs.

The JAK-STAT ligand Unpaired (Upd) is produced in specialized FCs called polar cells that are found at the anterior and posterior ends of developing egg chambers (Fig. 1A, D) (McGregor et al., 2002; Vied et al., 2012). Pathway activity is high in FCs in the germarium and decreases from posterior to anterior over the FSC domain with only low levels in ECs (Fig. 1A) (Vied et al., 2012). When JAK-STAT activity was elevated in FSC clones, it was shown that these FSCs out-competed wildtype FSCs and that proliferating FSC derivatives could accumulate in EC territory. Conversely, loss of STAT activity resulted in accelerated FSC loss (Vied et al., 2012). Those studies suggested potential roles in both FSC division and location or differentiation but detailed analysis was not possible at that time, before understanding of the organization and behavior of FSCs was drastically revised.

Here we have dissected cell autonomous responses to genetic changes in Wnt and JAK-STAT signaling pathways to separate their influences on each potentially independently controlled and separately measured parameter of FSC behavior: (1) FSC division rates, (2) FSC AP location, (3) FSC conversion to FCs, (4) FSC conversion to ECs, and (5) FSC competitive status measured by changes in FSC numbers over time (Fig. 1F). The results showed that the overall retention and amplification of FSCs can generally be explained in terms of changes in individual component behaviors, that these two graded pathways are substantially responsible for defining the FSC domain and that the dual role of the JAK-STAT pathway in promoting FSC division and FSC conversion to FCs contributes to co-ordination of FSC division and differentiation.

## Results

### Cell lineage approach to measure five separable parameters of FSC behavior

Prior to 2017, when each ovariole was thought to harbor just two or three FSCs, the cell autonomous effects of altered genotypes on FSC biology were ascertained by measuring the frequency of surviving marked FSC clones, defined by the presence of labeled FCs and a putative FSC, at various times after clone induction relative to control genotypes tested in parallel (Castanieto et al., 2014; Kirilly et al., 2005; O’Reilly et al., 2008; Song and Xie, 2003; Vied et al., 2012; Wang et al., 2012). Occasionally, the normally low frequency of ovarioles containing only marked FSCs and FCs (“all-marked”) was also elevated, indicating a genotype that increased FSC competitiveness. Now that it is appreciated that there are 14-16 FSCs in distinct AP locations, associated with different instantaneous division rates and differentiation potential, and that FSCs produce EC as well as FCs (Hayashi et al., 2020; Reilein et al., 2017), the results of clonal analysis can reveal far more about the effect of a specific genotype on FSC behavior. Correspondingly, labeled lineages must be scored in far more detail than before to reveal that information.

We conducted an extensive series of experiments using a standard regime in order to extract comprehensive information about FSC behavior and to be able to compare results for a large number of altered genotypes among all experiments in the series. We induced GFP-labeled clones in dividing cells of young adult females using the MARCM (Mosaic Analysis with a Repressible Cell Marker) system (Lee and Luo, 2001) with constitutive drivers (*actin-GAL4* and *tubulin-GAL4* together) of *UAS-GFP* and, where relevant, additional transgene expression. Heat-shock induction of a *hs-flp* recombinase transgene elicited recombination at the base of the relevant chromosome arm (using *FRT* recombination sites on 2L, 2R or 3R) in a fraction of FSCs (about 20%) to create homozygous recessive mutations or activate expression of a transgene (or both). After 6 or 12 days ovarioles were dissected, labeled for 1h with the nucleotide analog EdU to measure cell division (for the 6d test), fixed and stained to label all nuclei and the cell surface protein, Fasciclin 3 (Fas3). Each experiment included a variety of altered genotypes and a control with the same FRT recombination site. For each sample, GFP-labeled FC locations along the ovariole were recorded (Fig. 1C, D) and complete confocal z-section stacks of the germarium were archived and analyzed to count labeled FSCs in layer 1, immediately anterior to the border of strong Fas3 staining, and in the next two anterior layers (2 and 3), as well as labeled ECs (anterior to FSCs) (Fig. 1B), scoring also the number of labeled EC and FSCs that had incorporated EdU (at 6d) (Fig. 1E).

Our objective was to use the results to measure each distinguishable parameter of FSC behavior or decision-making separately (Fig. 1F). Cell division rates over the FSC domain were measured by EdU incorporation at the earlier time-point (6d) so that sufficient FSCs of poorly competitive genotypes were still present in good numbers and even hyper-competitive FSCs were not sufficiently abundant to potentially distort germarial morphology or induce secondary, non-autonomous responses. Genotype-dependent changes in the precise AP location of FSCs within the FSC domain were consistently most prominent at 12d, as were changes in the average number of FSCs present, so only those 12d results are presented. FC production was assessed quantitatively by a method we devised for measuring the probability of FC production per posterior FSC in one cycle of egg chamber budding (“p” in Fig. 1F), using 6d samples to ensure a suitably low frequency of posterior FSCs for all genotypes. EC production was measured from 0-6d and 0-12d; it was normalized to the inferred number of anterior FSCs present during those periods.

Normal FSC behavior was reported by controls from 31 separate MARCM experiments, scoring at least 50 ovarioles in almost every case. The results were extremely similar to those deduced previously from the more limited set of multicolor and MARCM experiments that formed the basis of our current perception of FSCs (Reilein et al., 2017). The results for controls are summarized in Fig. 1F and will be referenced individually later, in the context of genetic changes that alter those behaviors.

Here we note that each germarium contained an average of 3.2 marked FSCs at 6d and 3.3 at 12d, counting all ovarioles, including those with no labeled cells. If marked FSCs of the control genotype have no competitive advantage or disadvantage over unmarked FSCs it is expected that the average number of labeled FSCs should remain constant, as observed, and should therefore report the average number of FSCs initially labeled in each germarium. When deducing FSC properties for the first time it was often important to assay ovarioles with lineages derived from a single FSC (Fox et al., 2008; Kretzschmar and Watt, 2012; Reilein et al., 2017). In the present studies we already can define FSCs by location and the labeling of over three FSCs per ovariole is advantageous because it effectively allows us to examine the fate of a larger number of FSCs for a given number of ovarioles. Each FSC theoretically has a roughly equal and independent chance of being labeled by Flp-mediated recombination, so the number of labeled FSCs per ovariole can be estimated to have a binomial distribution centered around 3.2-3.3 (“0d”, red in Fig. 1G). The observed distribution at 6d was quite different from the assumed starting distribution, most obviously because more than a third of ovarioles no longer include any FSCs while the proportion of ovarioles with six or more FSCs has increased (Fig. 1G), consistent with expectations for neutral competition (Jones, 2010; Reilein et al., 2017; Reilein et al., 2018). These changes were further exaggerated at 12d but full colonization of a germarium by marked cells generally takes longer (Reilein et al., 2017), so even at 12d marked FSCs remain in a minority and are competing against unmarked wild-type cells in almost all ovarioles (Fig. 1G). That circumstance also applies to almost all variant genotypes investigated, ensuring all results reflect competition of marked FSCs with wild-type FSCs.

### Graded JAK-STAT signaling instructs graded FSC proliferation

In control clones 6d after induction, the average percentage of labeled FSCs that incorporated EdU was 25% (n = 4753) with a pronounced gradient of labeling, declining from posterior to anterior (33.4% for layer 1, 20.0% for layer 2, and 8.2% for layer 3) (Fig. 2A), similar to previous observations (Reilein et al., 2017). It had previously been observed that FSCs are rapidly lost in the absence of STAT activity, and that FSCs with excess JAK-STAT pathway activity became unusually numerous and incorporated EdU even within the EC domain, suggesting that this pathway may affect FSC proliferation (Vied et al., 2012). However, the quantitative effect of JAK-STAT signaling on FSC division rates has not been reported. We found that only 2.4% of FSCs with either of two homozygous null *stat* alleles incorporated EdU (n = 336), a ten-fold reduction compared to controls (Fig. 2A, B). To increase JAK-STAT activity, we expressed excess levels of the only Drosophila Janus Kinase, Hopscotch (Hop) using a *UAS-Hop* transgene in FSC clones (Vied et al., 2012; Xi et al., 2003), and found that 43.9% of UAS-Hop FSCs incorporated EdU (n = 1379), nearly double the rate of control FSC clones (Fig. 2A, C; Fig. S1). The pattern of proliferation in these FSCs still showed a posterior bias, with 49.9% of layer 1, 39.8% of layer 2, and 32.1% of layer 3 *UAS-Hop* FSCs incorporating EdU (Fig. 2A). These experiments demonstrated that the JAK-STAT pathway has a very strong positive, dose-responsive, cell autonomous influence on FSC division rate. We also saw that labeled cells in the EC region, which normally do not divide at all, sometimes incorporated EdU (12.4%) when JAK-STAT pathway activity was elevated (Fig. 2A; Fig. S1B), as noted previously without quantitation.

**Figure 2.**
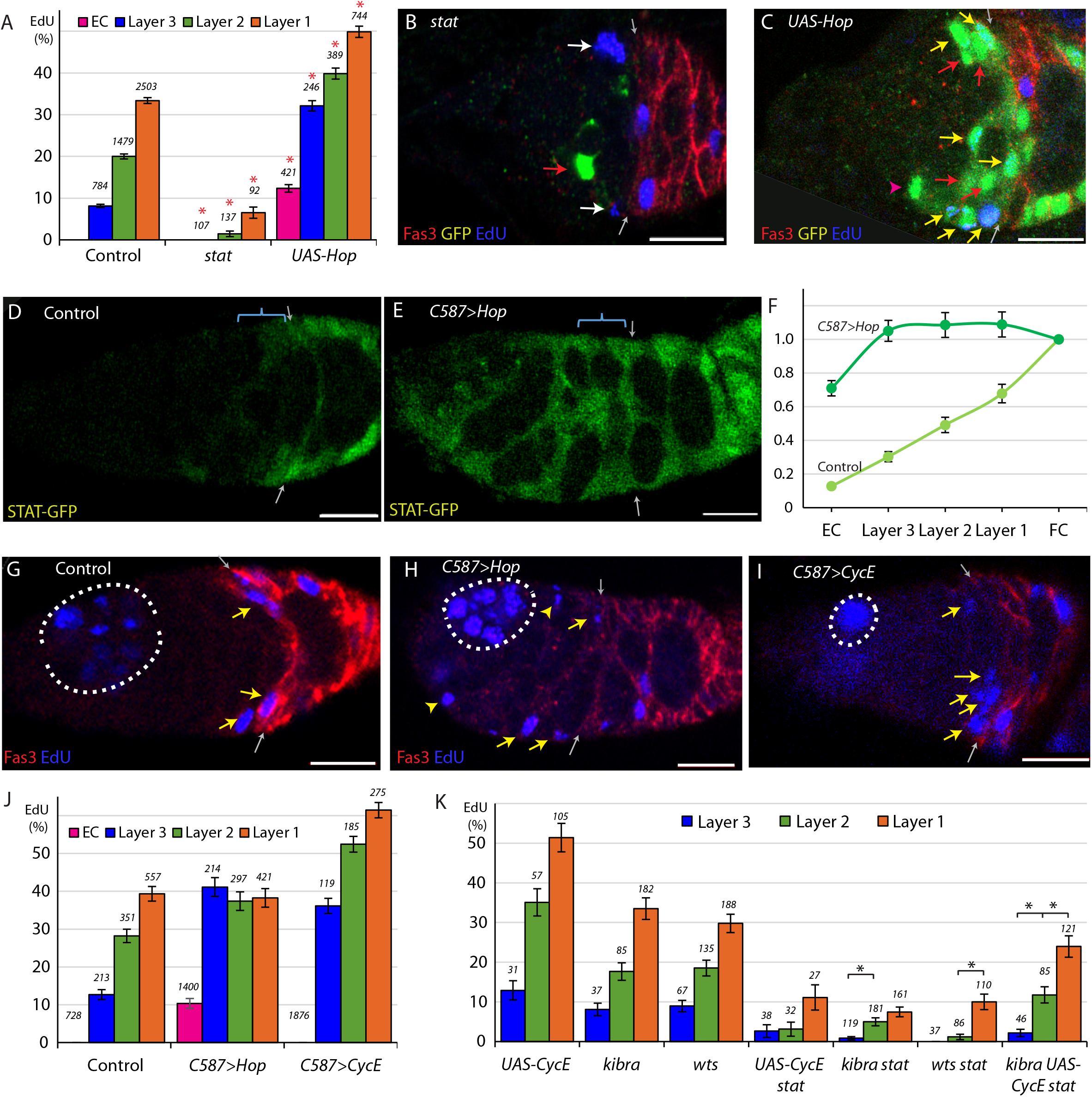
Graded JAK-STAT signaling determines FSC and EC proliferation profile. (A) EdU incorporation frequency into FSCs in layers 1-3 and ECs for the indicated genotypes of MARCM lineages, with the number of cells scored and significant differences to control values (red asterisks, p<0.001). (B, C) EdU (blue) incorporation into MARCM FSC lineages 6d after clone induction with anterior border of Fas3 (red) marked (gray arrows). (B) Most FSCs lacking *stat* activity (green) did not incorporate EdU (red arrow), unlike unmarked GFP-negative neighbors (white arrows). (C) Increased JAK-STAT pathway activity from expression of *UAS-Hop* produced many GFP-positive EdU^+^ FSCs (yellow arrows). GFP-positive EdU^−^ FSCs (red arrows) and a GFP-positive EdU^−^ EC (magenta arrowhead) are also indicated. (D-E) *STAT-GFP* reporter activity (green) (D) normally declines from the posterior over the FSC domain (blue bracket; arrows mark Fas3 (not shown) anterior border) but (E) becomes uniformly high 3d after UAS-Hop expression by C587-GAL4. (F) Average relative intensity of GFP fluorescence from STAT-GFP reporter in the indicated cell types for Control (n = 20) and *C587*>*Hop* (n= 22) germaria. (G-I) EdU (blue) incorporation in somatic cells is (G) normally restricted to FSCs (yellow arrows) and FCs beyond the Fas3 (red) border (gray arrows), (I) even when CycE activity is increased, but (H) ECs (arrowheads) are also labeled when JAK-STAT pathway activity is elevated. Dotted white lines outline germline cysts labeled by EdU. (J) EdU incorporation frequency into FSCs of layers 1-3 and ECs for germaria with the indicated genotypes (number of DAPI-labeled nuclei scored is above each column). (K) EdU incorporation frequency into FSCs of layers 1-3 and ECs for the indicated genotypes of MARCM lineages with number of cells scored above each column and significant differences between FSC layers indicated only for *stat*-containing genotypes (black asterisks, p<0.05). All scale bars are 10μm.

JAK-STAT pathway activity, reported by a “Stat-GFP” transgene with ten tandem STAT binding sites (Bach et al., 2007), is graded from posterior to anterior over the FSC domain (Fig. 2D, F), with a major ligand emanating from polar follicle cells (Vied et al., 2012). Because the JAK-STAT activity gradient runs parallel to the graded proliferation observed in EdU experiments we wished to test whether the two gradients were causally related. To do this, we took advantage of the *C587-GAL4* driver, which is expressed strongly in the anterior of the germarium and decreases in strength towards the posterior with no detectable expression in FCs (Reilein et al., 2017; Song et al., 2004). This pattern is roughly complementary to the JAK-STAT signaling pathway gradient. We expressed *UAS-Hop* from the *C587-GAL4* driver (*C587*>*Hop*), utilizing a temperature sensitive *GAL80* transgene (Zeidler et al., 2004) to restrict *UAS-Hop* expression temporally. After 3d at the restrictive temperature of 29C, we measured STAT-GFP fluorescence and found it to be similar in each of the three FSC layers and also over more anterior regions, indicating that the entire FSC and EC domains have roughly even JAK-STAT pathway activity (Fig. 2E, F).

With roughly normalized JAK-STAT activity across the FSC region, we measured proliferation in the three FSC layers. We observed nearly identical rates of division in each FSC layer; 42.8% of layer 1, 40.2% of layer 2, and 45.0% of layer 3 FSCs incorporated EdU in *C587*>*Hop* germaria (n = 932) (Fig. 2G, H, J). Thus, synthetically making JAK-STAT pathway activity uniform eliminated the normal posterior to anterior gradient of FSC division rates.

Additionally, elevated JAK-STAT activity in the anterior of the germarium stimulated EdU incorporation in 10.4% of ECs (Fig. 2H, J). In this experiment, those cells were quiescent ECs prior to increasing JAK-STAT activity. In the MARCM studies, the dividing cells in the EC region originated instead from labeled FSCs with elevated JAK-STAT signaling. Clearly, excess JAK-STAT pathway activity can stimulate cell division in the EC domain, whether the target cells are recently derived from FSCs or not. The rate of division of those cells was substantially lower than for cells in the FSC domain (Fig. 2J) despite similar levels of JAK-STAT pathway activity in the *C587-GAL4/UAS-Hop* experiment (Fig. 2F), suggesting the presence of other factors restricting EC division or inertia, perhaps resulting from the absence of anticipation of key cell cycle regulators in non-cycling cells (Spencer et al., 2013).

For comparison, we tested the effect of increasing CycE expression. In MARCM clones expressing *UAS-CycE* the FSC EdU index was greatly increased (Fig. 2K) but we observed no EdU incorporation in the EC region. When *UAS-CycE* was expressed with the *C587-GAL4* driver (*C587*>*CycE*) we found that the profile of EdU incorporation for FSCs remained graded as in control FSCs, contrasting with the response to UAS-Hop (Fig. 2I, J). Also, ECs remained quiescent. We conclude that graded JAK-STAT pathway activity instructs graded proliferation within the FSC domain, and that the anterior range of sufficient JAK-STAT pathway activity appears to define the anterior boundary of this critical stem cell property.

### JAK-STAT is not the sole potential contributor to the FSC proliferation gradient

If JAK-STAT signaling were uniquely responsible for regulating the FSC proliferation gradient, then we would not expect there to be any bias in EdU incorporation, by layer, for FSC clones that have no JAK-STAT pathway activity. Though EdU incorporation is very low in *stat* FSC clones, graded proliferation was still observed; 6.5% of layer 1 FSCs incorporated EdU, compared to 1.5% for layer 2 and 0% for layer 3 (Fig. 2A). Thus, there appear to be other influences that pattern FSC proliferation. Their magnitude is, however, hard to assess from this experiment alone.

We therefore sought to introduce additional genetic changes onto a *stat* mutant background. Crucially, those genetic changes should increase FSC proliferation without altering the normal graded pattern of proliferation. We found that expression of excess CycE and inactivation of upstream components of the Hpo pathway, Kibra and Warts (Wts), previously shown to influence FSC proliferation (Huang and Kalderon, 2014), fulfill this requirement (Fig. 2K). When each of these three manipulations was paired with *stat*, the average rate of proliferation roughly doubled compared to *stat* alone (2.4%), as 4.8% of *kibra stat* FSCs (n = 461), 5.2% of *wts stat* FSCs (n = 231), and 5.2% of *stat UAS-CycE* FSCs (n = 97) incorporated EdU, while *kibra* and *UAS-CycE* together increased EdU incorporation to 15.9% of *kibra UAS-CycE stat* FSCs (n=252) (Figure 2K). In all of these experiments, more FSCs in layer 1 incorporated EdU compared to the anterior layers in a pattern resembling that of normal FSCs (Fig. 2K), indicating that, in the absence of graded JAK-STAT pathway activity, there remains a robust mechanism for imposing graded FSC proliferation. Once the source of this mechanism is identified it will be possible to test whether it contributes to graded FSC proliferation under normal conditions or is effective only in the absence of JAK-STAT pathway activity.

### JAK-STAT pathway activity opposes anterior accumulation of FSCs and EC production

When scoring germaria with *stat* mutant clones, we observed that 96% of *stat* FSCs were found in the anterior layers of the germarium by 12d (Fig. 3A). We considered whether the reduced proliferation of *stat* FSCs may be contributing to this effect. For example, even though there is exchange of FSCs between layers, maintenance in layer 1 may depend on a particularly high rate of division because four times as many FSCs differentiate (to become FCs) in layer 1 than in the anterior layers (to become ECs) in each budding cycle (Reilein et al., 2018). When we tested other mutants that had severely impaired proliferation (Wang et al., 2012; Wang and Kalderon, 2009), we did observe a small reduction in the proportion of FSCs in layer 1 from a control value of 49% to 39% for *cycE* and 43% for *cutlet* FSCs but the change was much smaller than observed for *stat* mutant FSCs (Fig. 3A).

**Figure 3.**
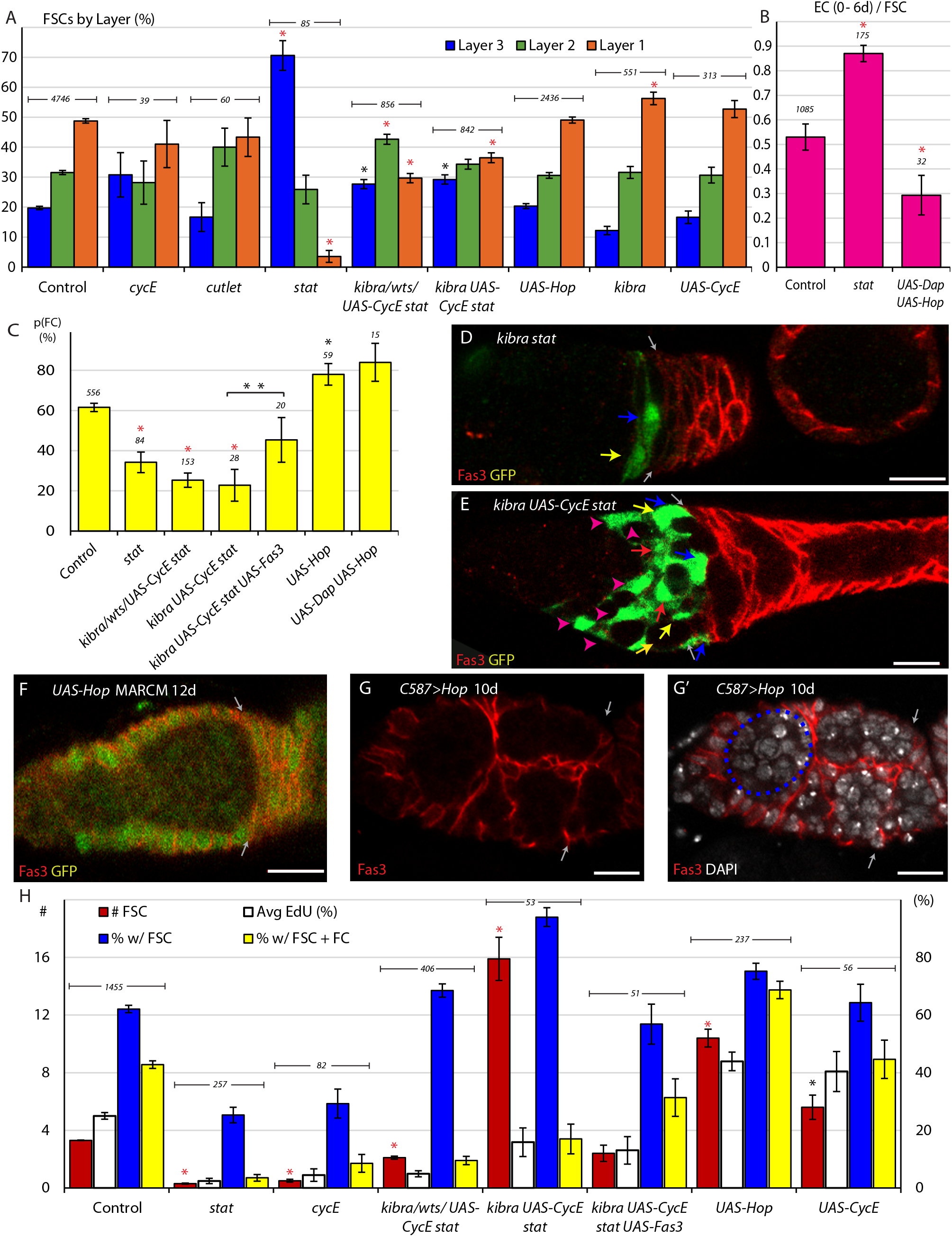
JAK-STAT promotes conversion of FSCs to FCs. (A) Relative frequency of marked FSCs in each layer for the indicated genotypes of MARCM lineages, with the number of total FSCs scored for each genotype and significant differences to control values (black asterisks, p<0.05, red asterisks, p<0.001). (B) ECs produced per anterior FSC from 0-6d for the indicated MARCM lineage genotypes with the total number of relevant germaria scored and significant differences to control values (red asterisks, p<0.001). (C) Average probability of a layer 1 FSC becoming an FC during a single budding cycle for the indicated MARCM lineage genotypes with the number of informative germaria scored and significant differences to control values or for the bracketed comparison showing the impact of *UAS-Fas3* (black asterisks, p<0.05, red asterisks, p<0.001). (D-E) Despite marked FSCs in layer 1 (blue arrows), and a significant number of layer 2 FSCs (yellow arrows), layer 3 FSCs (red arrows), and ECs (magenta arrowheads), marked FCs, posterior to the Fas3 (red) border (gray arrows) are absent here (and are generally rare) in *kibra stat* and *kibra UAS-CycE stat* MARCM lineages. (F-G) Increased JAK-STAT pathway activity induced ectopic anterior Fas3 (red) expression (F) cell autonomously in MARCM lineages (green) and (G-G’) in numerous cells anterior to the normal Fas3 border (arrows) when increased throughout the anterior germarium using C587-GAL4, sometimes partitioning single cysts, visualized by DAPI (white) nuclear staining, into egg chamber-like structures (dashed blue line). (H) Number of FSCs per germarium (red), frequency of FSCs incorporating EdU (aggregating all layers, white), frequency of ovarioles with a marked FSC (blue) and frequency of ovarioles with a marked FSC and marked FCs (yellow) for the indicated genotypes, with the number of germaria scored at 12d (EdU was scored at 6d with n reported in (A)) and significant differences for FSC numbers compared to control values (black asterisks, p<0.05, red asterisks, p<0.001). All scale bars are 10μm.

We also tested the consequences of increasing the division rate of *stat* mutant FSCs using the *kibra, wts,* and *UAS-CycE* manipulations. The proportion of layer 1 FSCs was (on average) 29.7% for *kibra stat, wts stat*, and *UAS-CycE stat* FSCs, and 36.5% for *kibra UAS-CycE stat* FSCs, still much lower than for controls (49%) (Fig. 3A, D, E). Thus, we observed a consistent anterior bias for all FSCs lacking STAT activity, even for genotypes that permitted division at rates approaching normal values. The observation that *stat* mutant FSCs with higher division rates had a reduced anterior bias also support the hypothesis that reduced division rates selectively deplete FSCs from the fastest-dividing posterior layer.

The average number of marked ECs accumulated by 6d was divided by the average number of anterior marked FSCs present at 6d to measure EC production per anterior FSC over 0-6d as 0.53 for control clones. The calculation for altered genotypes uses measurements of control and mutant anterior FSC numbers at 6d to estimate the average number of anterior mutant FSCs over the 0-6d period (see Methods); it revealed an increase in the number of ECs produced per anterior FSC in the absence of STAT activity (0.87) (Fig. 3B). Thus, loss of STAT activity resulted in more anterior locations for FSCs and a greater frequency of conversion of anterior FSCs to ECs, their anterior neighbors.

Increased JAK-STAT pathway activity did not significantly alter the location of FSCs or EC production, tested with *UAS-Hop* alone or together with *UAS-Dap* (*Dacapo*) (Lane et al., 1996; Lehner et al., 1992) to moderate effects on FSC division rate (Fig. 3A, B). These results suggest that a certain (unknown) minimal level of pathway activity is required to prevent unbalanced anterior migration of FSCs and accelerated conversion of anterior FSCs to ECs.

### JAK- STAT pathway activity promotes FC production from posterior FSCs; role of Fas3

The signals and mechanisms that govern conversion of an FSC to an FC are largely unknown. In fact, only with recent insights into FSC organization can we measure this process independently of other factors, such as altered division rates, in order to attribute changes in FSC survival to changes in the frequency of differentiation to FCs (Fig. 1F). Importantly, an FSC can become an FC at any time relative to its last division, and FSC division and differentiation can therefore potentially be regulated independently (Reilein et al., 2018). Previous studies also found that a single founder FC produced a patch occupying 17.8% of the monolayer of an egg chamber on average, which translates to an average of 5.6 founder FCs (1/0.178) produced per cycle of egg chamber budding (Reilein et al., 2018).

We devised a method to determine the probability that a single FSC becomes an FC from our MARCM data. We can recognize FCs produced in the latest cycle as being in the first layer of Fas3-positive cells (“Immediate FCs”; Fig. 1B-D); we can easily score if there are none but we cannot reliably score their number. We therefore scored the presence or absence of Immediate FCs in germaria with 0-3 marked posterior FSCs in 6d samples. Germaria with higher numbers of marked FSCs almost invariably include marked immediate FCs and are not informative. The calculations factored in measured differences in FSC division rate for different genotypes because that influences the total number of FSCs available for conversion to FCs during a cycle (see Methods). For 556 control germaria across 31 MARCM experiments, we found that a layer 1 FSC has, on average, a 61.6% likelihood (p=0.616) of becoming an FC in one cycle (Fig. 3C). Under the assumptions of this method, the average number of layer 1 FSCs present during a cycle was 9.93 (7.6 plus (7.6× 7/16 × ½) new FSCs), so that 9.93 × 0.616 = 6.1 FCs would be produced per cycle according to the experimentally determined p value. The result is close to the value of 5.6 calculated from founder FC clone sizes, validating the method employed to calculate the probability of FC formation per FSC.

Using the same method, we found that a *stat* mutant layer 1 FSC was much less likely than controls (34.2% compared to 61.6%) to produce an FC (Fig. 3C). However, as we previously noted, the majority of *stat* FSCs do not populate layer 1 and the sample size was consequently small, even over several experiments. We then looked at *kibra stat, wts stat, and UAS-CycE* together with loss of STAT, which all had a greater number of FSCs in layer 1. In these samples, we found the average likelihood of an FSC becoming an FC to be 25.3%, and 22.8% for *kibra UAS-CycE stat* mutant FSCs (Fig. 3C, D). These experiments demonstrated that FC production from its immediate precursor, a layer 1 FSC is significantly impaired in the absence of STAT activity.

We also examined the consequences of increasing JAK-STAT pathway activity in a layer 1 FSC. We found that the probability of becoming an FC increased to 78.1% for FSCs expressing *UAS-Hop* (Fig. 3C). Here the sample size was relatively small (12 germaria per experiment for a total of 59 germaria) because only germaria with few FSCs are informative and FSCs with excess JAK-STAT pathway are quite abundant even by 6d. We therefore introduced a *UAS-Dacapo* (*UAS-Dap*) transgene, encoding a CycE/Cdk2 inhibitor (Lane et al., 1996; Lehner et al., 1992), to limit proliferation rates. We found that 23.8% of *UAS-Dap UAS-Hop* FSCs incorporated EdU, a rate similar to controls. Layer 1 FSCs expressing both *UAS-Hop* and *UAS-Dap* had a significantly higher probability (84.0%) than controls of becoming FCs (Fig. 3C). Thus, both increased and decreased JAK-STAT pathway activity significantly affected the production of FCs from FSCs, suggesting that the magnitude of pathway activity is an important factor in regulating this transition.

The conversion frequency of layer 1 FSCs to FCs was not greatly altered by increased CycE expression (71%); it was decreased substantially for *cycE* (35%) and *cutlet* (35%) mutant FSCs but not for *yki* (51%) or *smo* (74%) mutant FSCs, all of which greatly reduce FSC division rates (Huang and Kalderon, 2014; Wang et al., 2012), and was not restored for *cutlet* FSCs expressing *UAS-CycE* (39%), which divide faster than controls. Thus, it remains uncertain whether FSC division rate *per se* might affect layer 1 FSC to FC conversion. However, the pronounced effects of altering JAK-STAT activity on altering FC production probability are not secondary to changes in division because equally large changes were observed over a variety of FSC division rates for both decreased and increased pathway activity.

The proportion of FSCs in layer 1 depends not only on movements between FSC layers but also on the rate of depletion from layer 1 to form FCs. Loss of STAT activity was found to reduce layer 1 occupancy even though conversion of layer 1 FSCs to FCs was reduced, suggesting that the bias towards anterior movement within the FSC domain is even stronger than measured simply by steady-state AP distribution (Fig. 3A). Similarly, excess JAK-STAT pathway did not affect steady-state AP location but increased conversion of layer 1 FSCs to FCs, suggesting that there is in fact a bias towards posterior movement within the FSC domain.

It was previously noted that strong expression of the surface adhesion molecule Fas3, which is normally observed only in FCs, was seen in some derivatives of FSCs with elevated JAK-STAT signaling in the EC and FSC domains (Vied et al., 2012). We confirmed these observations for MARCM clones, finding that 66% of germaria with labeled cells in the anterior half of the germarium (the FSC and EC domains) showed ectopic Fas3 expression at 12d after clone induction (Fig. 3F). Furthermore, we observed that these cells sometimes appeared to form a crude epithelial monolayer surrounding developing germline cysts, indicative of FC behavior. Similar structures were observed in germaria where *UAS-Hop* was conditionally expressed using *C587-GAL4* (Fig. 3G). Here, 53% of germaria expressed ectopic Fas3 by 3d, increasing to 72% by 6d and 94% by 10d. Thus, high JAK-STAT pathway alone can instruct at least some aspects of the FC phenotype even in locations where FCs do not normally form.

To test whether Fas3 might be an important intermediate in the normal influence of JAK-STAT signaling on FC production we expressed excess Fas3 in *kibra UAS-CycE stat* mutant FSCs. We observed a doubling in layer 1 FSC to FC conversion (from 22.8% to 45.4%) (Fig. 3C), suggesting that increased Fas3 expression can partially restore FC production in the absence of JAK-STAT pathway activity.

### Net effect of JAK-STAT on cell-autonomous FSC longevity and amplification

By measuring the impact of altered genotypes on each component of FSC behavior (Fig. 1F) it should be possible to predict, or at least rationalize, the net effect on FSC competitive behavior in MARCM lineage analyses, measured by the proportion of ovarioles that retain marked FSCs over time, or more precisely by the average number of marked FSCs over all ovarioles.

For FSCs and other stem cells governed by population asymmetry in which differentiation is independent of stem cell division, the rate of stem cell division is necessarily a major determinant of competitive success (Reilein et al., 2018). Accordingly, by 12d, there were an average of 10.4 *UAS-Hop* FSCs per germarium, significantly greater than the 3.3 FSCs per germarium observed in controls (Fig. 3H). The proportion of ovarioles containing a marked FSC was also increased with *UAS-Hop* expression to 75.2% compared to 62.1% in controls (Fig. 3H). FSC clones expressing *UAS-CycE* have a similar increase in division rate to those expressing *UAS-Hop* (Fig. 2A, K), as measured by average EdU index (40.4% vs 43.9%) but only yielded an average of 5.6 marked FSCs per ovariole by 12d with 64.3% of ovarioles containing a marked FSC (Fig. 3H). Neither *UAS-CycE* nor *UAS-Hop* significantly altered steady-state FSC AP location (Fig. 3A), and while *UAS-Hop* promoted conversion of FSCs to FCs, *UAS-CycE* did not. The larger impact of increased JAK-STAT pathway activity on FSC numbers is plausibly because increased division was promoted preferentially in anterior FSC layers (Fig. 2A, K), which normally do not divide as frequently and are lost directly to differentiation at a lower frequency than posterior FSCs.

When STAT activity was eliminated in FSC clones, there were an average of 0.3 *stat* FSCs per germarium and 25.3% of germaria retained a marked FSC after 12d (Fig. 3H). These are large deficits compared to controls (3.2 FSCs, 62.1% of ovarioles), and similar to *cycE* mutants (0.5 FSCs, 29% of ovarioles), which also drastically reduce FSC proliferation. When we tested *kibra stat, wts stat,* and *UAS-CycE stat* genotypes, the average number of FSCs increased to 2.1 per germarium with 68.5% of germaria containing an FSC clone. An improved persistence of FSCs was expected but the magnitude of rescue was surprisingly large, given that these FSCs still divided at only one fifth the rate of control FSCs. This disparity was even more pronounced when examining FSC competition for *kibra stat* mutants expressing *UAS-CycE*, which divided at about 64% of wild-type rates. Here, the average number of marked FSCs was 15.9 and 94% of ovarioles included marked FSCs (Fig. 3H). The remarkable persistence and amplification of these FSCs likely results from the infrequent presence of FSCs lacking STAT activity in layer 1 (Fig. 3A) and the markedly lower conversion of posterior FSCs into FCs (Fig. 3C). The virtual sealing off of this conduit greatly outweighs a small increase in conversion to ECs because the normal rate of FSC conversion to ECs is four-fold lower than conversion to FCs.

Although the survival and amplification of FSCs lacking STAT activity can be increased towards, and then beyond normal by relatively modest restoration of division rates, those FSCs still show much reduced physiological activity, measured by continued production of FCs, with very few ovarioles containing both FSCs and FCs (3.5% for *stat*, 9.5% for *stat* with *kibra*, *wts* or *UAS-CycE*, 17.0% for *kibra UAS-CycE stat*, compared to 42.8% for controls) (Fig. 3D, E, H). Thus, interfering with the normal coordination of FSC division and conversion to FCs in response to JAK-STAT pathway activity by adding genetic modifiers of division alone can lead to extensive amplification of unproductive FSCs.

When excess Fas3 was expressed in *kibra UAS-CycE stat* FSCs, doubling conversion of layer 1 FSCs to FCs, the average number of FSCs per germarium declined sharply from 15.9 to 2.4 (Fig. 3H). At the same time, the percentage of ovarioles with at least one FSC at 12d declined from 94% to 59% but the proportion of ovarioles with FSCs and FCs increased from 17% to 31% (Fig. 3H). The response to excess Fas3 emphasizes the large impact of the rate of FC production on FSC numbers and their ability to fulfill their physiological role. It also demonstrates that appropriate magnitudes of artificial stimulation of both FSC division and FSC differentiation to FCs can partially substitute for the normal coordination of these rates by JAK-STAT signaling to bring about roughly normal FSC behavior.

### Wnt signaling primarily influences FSC position and differentiation

The role of Wnt signaling in regulating FSC behavior has already been examined in the context of a revised model of FSC numbers, locations and properties. A *Fz3-RFP* reporter demonstrated that Wnt pathway activity decreases in strength across the FSC region, in the anterior to posterior direction (Reilein et al., 2017; Wang and Page-McCaw, 2014). FSCs with an arrow (*arr*) mutation to eliminate the Wnt pathway response and axin *(axn)* or Adenomatous Polyposis Coli (*apc*) mutations to constitutively activate the Wnt pathway in MARCM clones (Reilein et al., 2017), illustrated a consistent effect of higher Wnt signaling activity favoring anterior FSC locations and greater conversion to ECs; 77.6% of *arr* FSCs but only 15-20% of *axn* and *apc* FSCs were observed in layer 1, while 9.1 *axn* and *apc* ECs and 0.1 *arr* ECs were observed per germarium, compared to an average of 1.5 ECs for controls (Reilein et al., 2017).

Here we tested the effect of reducing rather than eliminating Wnt pathway activity by expression of a *UAS-dnTCF* transgene (van de Wetering et al., 2002) in clones. We found that 63.0% of *UAS-dnTCF* FSCs were observed in layer 1 (compared to 48.8% for controls), a significant change but less pronounced than for *arr* FSCs, which showed a layer 1 occupancy of 79.3% across three replicates (n = 237 cells), including two additional tests not previously reported (Fig. 4A). We also tested an additional *axn* replicate and confirmed layer 1 occupancy to be greatly decreased, to 20.6% at 12d. Layer 1 occupancy was slightly higher (31.0%) for *axn* FSCs that expressed excess CycE (Fig. 4A) to increase division rates (from 7.4% to 15.3% towards control 25.0% EdU frequency (Fig. 4E)), consistent with earlier evidence of reduced division favoring more anterior locations.

**Figure 4.**
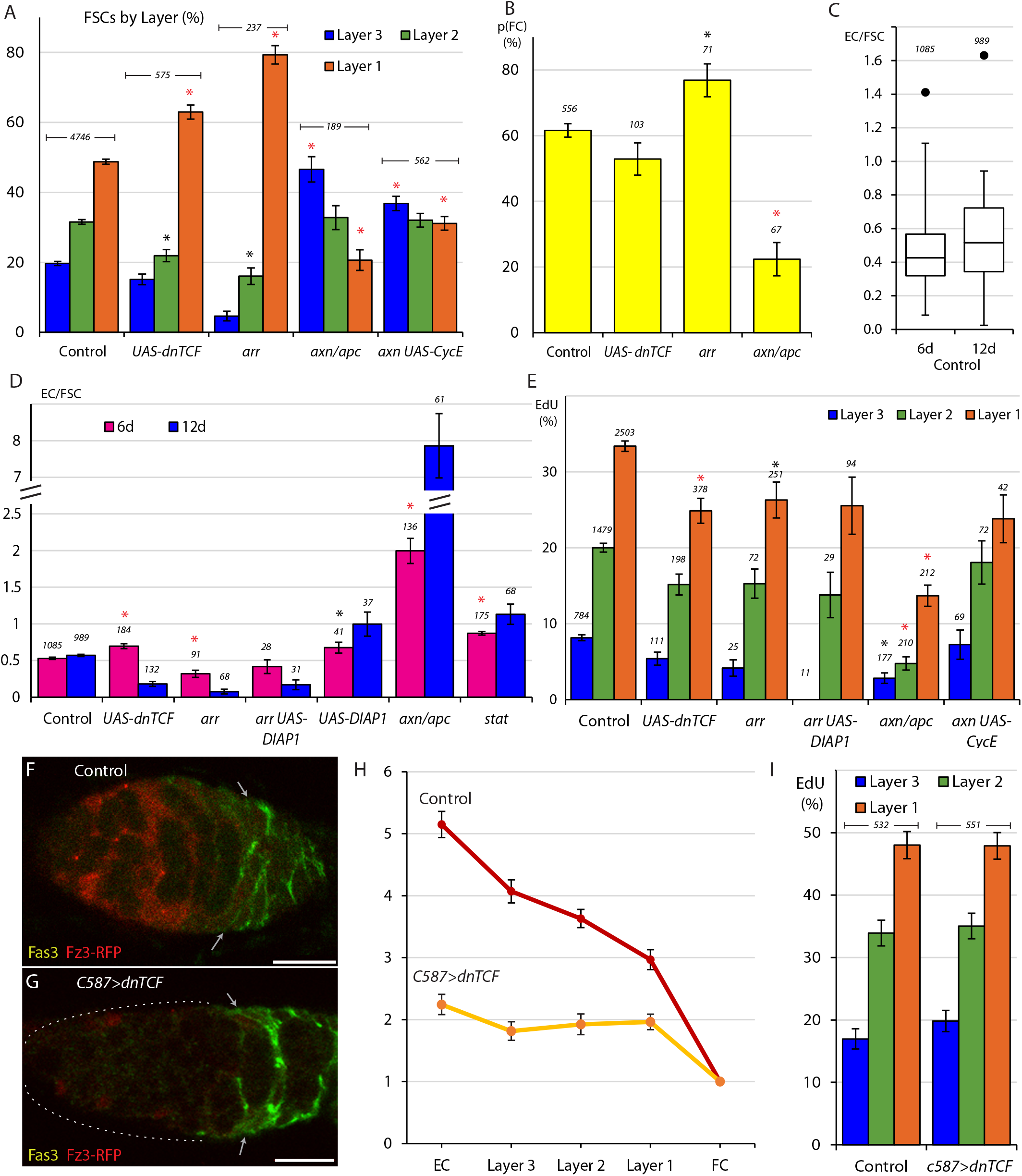
Wnt signaling opposes FC production and promotes anterior FSC location and EC production. (A) Relative frequency of marked FSCs in each layer for the indicated genotypes of MARCM lineages, with the number of total FSCs scored for each genotype and significant differences to control values (black asterisks, p<0.05, red asterisks, p<0.001). (B) Average probability of a layer 1 FSC becoming an FC during a single budding cycle for the indicated MARCM lineage genotypes with the number of informative germaria scored and significant differences to control values (black asterisks, p<0.05, red asterisks, p<0.001). (C) Box-and-whisker plot of ECs produced per anterior FSC from 0-6d and 0-12d across all MARCM controls (n = 31 experiments) with a single outlier and the number of germaria scored. (D) ECs produced per anterior FSC from 0-6d (magenta) and 0-12d (blue) for the indicated MARCM lineage genotypes with the total number of informative germaria scored and significant differences to control values (black asterisks, p<0.05, red asterisks, p<0.001). (E) EdU incorporation frequency into FSCs of layers 1-3 and ECs for the indicated genotypes of MARCM lineages with number of cells scored above each column and significant differences between FSC layers indicated only for *stat*-containing genotypes (black asterisks, p<0.05, red asterisks, p<0.001). (F-G) Fz3-RFP reporter of Wnt pathway activity (red) (F) normally declines in strength from anterior to posterior but (G) was mostly eliminated after 3d of *UAS-dnTCF* expression with *C587-GAL4*. (H) Average Fz3-RFP intensity for control (n = 23 germaria) and C587>dnTCF (n = 22 germaria) genotypes. (I) EdU incorporation frequency into FSCs in layers 1-3 for the indicated genotypes (number of DAPI-labeled nuclei scored is above each column). All scale bars are 10μm.

We also used the “Immediate FC” method to measure FC production. We found that *arr* layer 1 FSC clones have an elevated probability (76.9% compared to 61.6% in controls) of becoming an FC, (Fig. 4B). That response was not reproduced by reducing Wnt pathway activity with *UAS-dnTCF* (52.9% probability). We also observed that FSCs with increased Wnt activity showed only a 22.4% likelihood of becoming an FC, a roughly threefold decrease from control values (Fig. 4B). We therefore extend previous conclusions to surmise that the AP location of FSCs in a competitive environment of normal FSCs is altered by reduction, elimination or increases of Wnt pathway activity. Increased pathway activity also reduces FC production from posterior FSCs but only the most severe reductions in Wnt pathway activity enhance FC production. These results suggest that the magnitude of Wnt pathway activity affects AP migration over the whole FSC domain, where Wnt signaling is graded, and that the decline to near zero values at the posterior margin of the FSC domain is a significant determinant of the FSC to FC transition.

### EC turnover rate is influenced by Wnt and JAK-STAT pathway activities

To evaluate EC production from anterior FSCs, we calculated the ratio of ECs per anterior FSC, in germaria that retained at least one marked FSC. In controls, we found this ratio was 0.53 (SE 0.30; SEM 0.05) for the period from 0-6d and 0.57 (SE 0.29; SEM 0.05) for the period from 0-12d (Fig. 4C). If ECs are produced by FSCs at a constant rate and all labeled ECs accumulate without loss, we would expect this ratio to double from 6d to 12d, reflecting a longer period of accumulation from a roughly constant supply of labeled anterior FSCs. The observed percentage increase was much lower (8%) across 31 control tests. Thus, there must be considerable EC turnover, leading to the inferred loss of more than half of the marked ECs in the 6d following their production from 0-6d. Apoptosis is observed in normal germaria at a low frequency (Pritchett et al., 2009). Likewise, it is plausible that an EC can occasionally move into FSC territory. However, if all of the roughly forty ECs behaved like marked ECs produced from adult FSCs, experiments that examined sole FSC candidates in FSC lineages would have revealed many cells in the EC region to be stem cells. Likewise, observed cell death rates cannot account for such a high turnover of all ECs. We therefore deduce that ECs produced during adulthood are less stable than those present at eclosion.

By altering FSC genotypes we might hope to distinguish between EC death and reversion to FSC status and to measure the effects of signaling pathways on EC turnover as well as EC production. However, the variability in these measures among controls (Fig. 4C) indicates that observed changes must be robust to be significant indicators of altered behavior. We were not yet able to make a reliable determination of the role of apoptosis by expressing the inhibitor of apoptosis, DIAP1 in otherwise wild-type clones. In a single experiment, we found slightly elevated EC production from 0-6d (0.68) and a greater increase for DIAP1-expressing FSCs from 0-12d (1.0), indicating slower turnover but still short of the expected doubling for completely stable ECs (Fig. 4D).

We found that EC production from 0-6d was reduced for *arr* mutant FSCs (0.32 compared to 0.53 for controls) but not for *UAS-dnTCF* FSCs (0.70). However, from 0-12d EC production was even lower for both genotypes (0.08 for *arr* and 0.18 for *UAS-dnTCF*), representing a decline (net EC loss) of 24% (*arr*) and 59% (*UAS-dnTCF*) from 6-12d (Fig. 4D). These data indicated that loss or reduction of Wnt signaling greatly increased the turnover of ECs derived from FSCs during adulthood.

Complete loss of Wnt pathway activity is known to elicit apoptosis in ECs, especially in anterior locations (Wang et al., 2015; Wang and Page-McCaw, 2018). A single experiment showed that when *arr* mutant FSCs expressed *DIAP1* EC production from 0-6d was increased (0.42) relative to *arr* alone (0.08) but EC production from 0-12d was still lower (0.17), suggesting net loss of labeled ECs from 6-12d, and contrasting with the findings from expression of DIAP1 alone (a 48% increase) (Fig. 4D). The implication, pending further testing, is that loss of Wnt signaling may accelerate reversion of ECs to FSCs.

When Wnt pathway activity was increased using the *axn* and *apc* mutations we found the average ratio of ECs per anterior FSC to be 2.0 from 0-6d and 7.86 from 0-12d, revealing an increase of over 300% from 6-12d (Fig. 4D). This indicates accelerated EC production over time and presumably, little or no turnover of newly-produced ECs, suggesting that any normal reversion of ECs to FSCs is strongly opposed by high levels of Wnt pathway activity.

We also investigated JAK-STAT signaling and EC production dynamics. When STAT activity was eliminated, we found that EC production increased to 0.87 EC/FSC from 0-6d and 1.1 EC/FSC from 0-12d (Fig. 4D), showing an average 42.2% EC increase from 6-12d, a slightly higher rate than in controls, suggesting the possibility that EC to FSC reversion is reduced by loss of STAT activity.

### Wnt signaling can reduce FSC division but does not normally affect the magnitude or pattern of FSC division significantly

It has previously been reported that loss of Wnt signaling reduced FSC division by a small amount, with most measured FSCs in layer 1, while increased Wnt pathway activity, measured mostly in anterior FSCs, greatly decreased FSC proliferation, with results normalized in each case for FSC locations (Reilein et al., 2017). To examine the effects of reduced signaling more comprehensively in anterior FSCs we looked at more *arr* mutant samples, including those additionally expressing DIAP1, and four independent experiments where Wnt signaling was reduced by expression of *dnTCF*, which results in a less pronounced posterior shift of FSCs than complete inhibition of Wnt signaling. We found that EdU incorporation was marginally reduced in all layers for FSCs lacking *arr* activity (26.3%, 15.3%, 4.2% for layers 1-3) or expressing *dnTCF* (24.9%, 15.2%, 5.4% for layers 1-3) relative to controls (33.4%, 20.0%, 8.2% for layers 1-3). In all experiments where Wnt signaling was reduced or eliminated there was a clear posterior to anterior proliferation gradient, as in normal FSCs (Fig. 4E).

In response to increased Wnt pathway activity, the EdU index was substantially reduced in all layers (13.7%, 4.8%, 2.8% for layers 1-3) but a posterior to anterior gradient was still evident (Fig. 4E). To test the effect on graded proliferation further we combined loss of *axn* with *UAS-CycE* in an attempt to restore overall FSC proliferation towards wild-type levels. Excess CycE indeed doubled division rates overall (from 7.4% to 15.3% EdU incorporation; control was 25%) and a robust posterior to anterior gradient was still evident (23.8%, 18.1%, 7.3% for layers 1-3) (Fig. 4E).

Finally, to assess the contribution of graded Wnt signaling to the A/P pattern of graded proliferation, we reduced Wnt signaling globally by expressing the *UAS-dnTCF* transgene with the *C587-GAL4* driver (*C587*>*dnTCF*). As both normal Wnt pathway activity and *C587-GAL4* expression decline from anterior to posterior across the FSC domain, this manipulation ought in theory to flatten or eliminate the normal Wnt gradient to produce a roughly even, low level of pathway activity. Fz3-RFP reporter expression showed that Wnt signaling was considerably reduced and close to uniform across the three FSC layers (Fig. 4F-H). Under these conditions there was very little difference in EdU incorporation in any FSC layer when compared to controls (Fig. 4I). This result suggests that graded FSC proliferation does not rely on graded Wnt signaling activity even though increases in Wnt pathway activity have the potential to inhibit FSC division.

### JAK-STAT and Wnt pathways do not directly (cell autonomously) affect one another

As both the JAK-STAT and Wnt signaling pathways play important roles in the regulation of FSCs, we asked whether regulation by each pathway is accomplished independently. We first measured whether genetic manipulations influenced Wnt pathway activity by inducing GFP-positive MARCM clones that also expressed the Fz3-RFP reporter. In these tests, we measured the signal intensity of the reporters in the marked cells and unmarked neighbors (Fig. S2). Cells were considered neighbors if they would have been scored in the same FSC layer, and if they were captured within the same z-section during confocal imaging to ensure a consistent signal intensity. Samples were examined 6d after clone induction so that all genotypes included marked cells in a full range of FSC locations.

For genotypes affecting Wnt signaling components, Fz3-RFP intensity was significantly altered when compared to neighbors; reporter activity was significantly different (p=0.001) at 39.7% in *arr* clones compared to neighbors and 63.1% for marked FSCs expressing *UAS-dnTCF* clones (p=0.01) (Fig. S2). A null *arr* genotype is expected to prevent all stimulated Wnt pathway activity, so the measured residual RFP likely corresponds principally to background staining and perduring RFP expressed before Arr protein was fully depleted during clone expansion. *UAS-dnTCF* therefore likely reduces Wnt pathway activity to below 40% of normal levels (23.4% reduction in a range of 60.3% from null to normal) in MARCM FSC lineages. By the same logic, the measured elevation of Fz3-RFP levels in *axn* clones (169.4%, p=0.02) corresponds to a roughly two-fold increase in normal pathway activity (129.5% change compared to a normal range of 60.3%) (Fig. S2).

FSCs with no STAT activity had, on average, 108.4% of neighboring cell Fz3-RFP intensity (p=0.74), while FSCs expressing *UAS-Hop* had 84.8% of control cell levels (p=0.41). Additionally, we tested *kibra UAS-CycE stat* clones (103.5%) and found no significant change (p=0.74) in Fz3-RFP intensity (Fig. S2C). Thus, altering JAK-STAT pathway activity did not have a major direct, cell autonomous influence on Wnt pathway activity.

We then asked how manipulating both pathways within the same FSC would influence its behavior, and therefore assessed measurements of proliferation, position, and differentiation when both JAK-STAT and Wnt signaling activity were altered in FSC clones.

### FSC division rates; inhibition by Wnt is suppressed by JAK-STAT pathway activity

Reduction or elimination of Wnt signaling alone barely affected FSC proliferation, producing a small reduction at most. It was therefore surprising that expressing *UAS-dnTCF* increased EdU incorporation in FSCs lacking STAT activity (from 2.4% to 4.1%). This effect was more striking in STAT-deficient genotypes that increased division rate, elevating the EdU index of FSCs with *stat* mutations together with *kibra, wts,* or *UAS-CycE* from 5.0% to 16.2% (Fig. 5A), close to the division rates observed for otherwise normal FSCs expressing dn-TCF (19.0%). Two separable contributions of dnTCF can be resolved in these latter genotypes. First, dnTCF increased the proportion of marked FSCs in layer 1 from 27.6% to an intermediate value (36.9%), still lower than for controls (48.4%) or expression of dnTCF alone (60.3%) (Fig. S4D). Second, and more importantly, the EdU index was increased in each layer (22.6% vs 8.7%, 14.3% vs 5.7%, 5.5% vs 1.0% in layers 1-3, respectively) (Fig. 5A).

**Figure 5.**
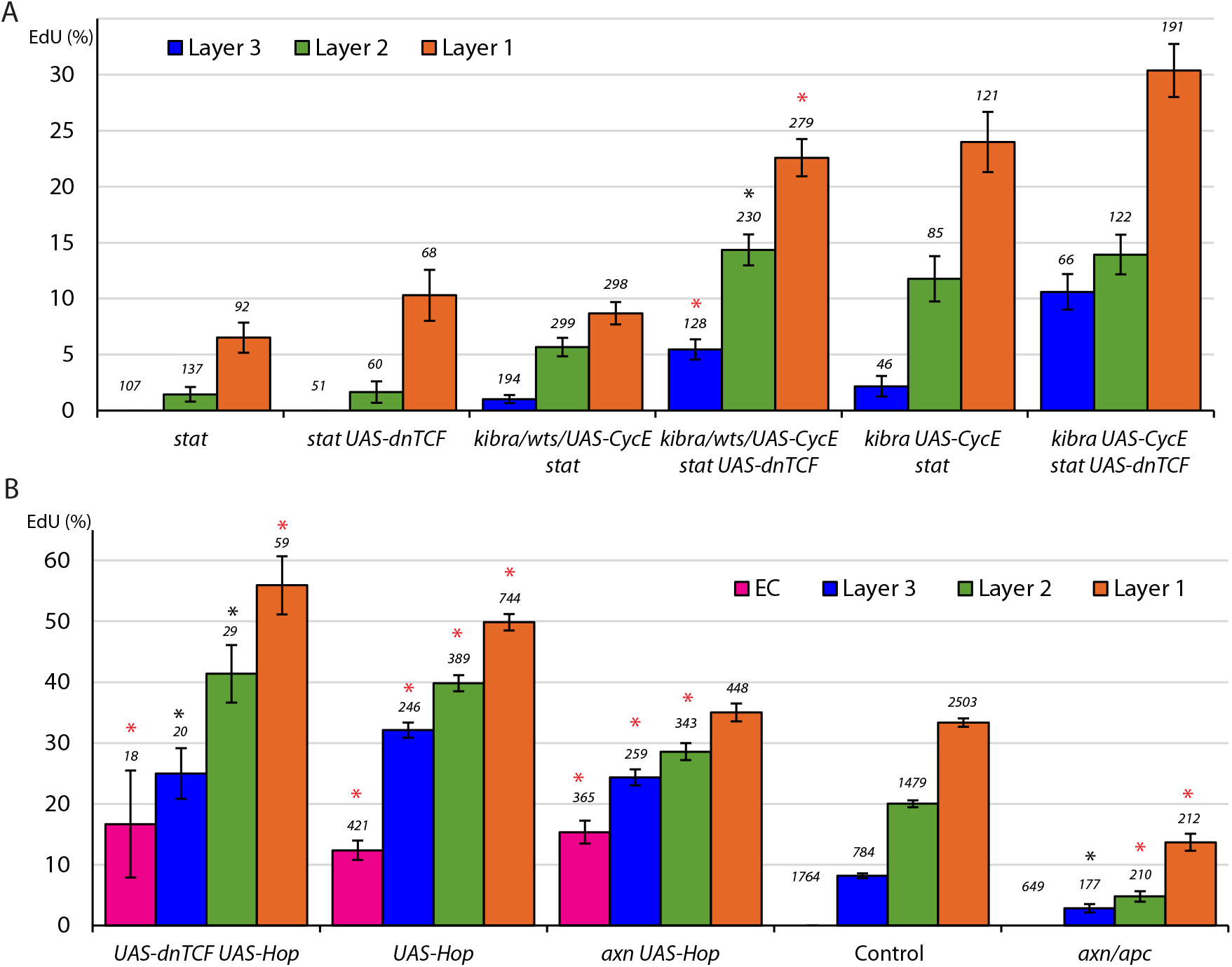
Wnt pathway activity reduces FSC division only in the absence of JAK-STAT pathway activity and JAK-STAT overrides inhibition by the Wnt pathway when both are in excess. (A, B) EdU incorporation frequency into FSCs of layers 1-3 and ECs for the indicated genotypes of MARCM lineages with number of cells scored above each column and (A) significant differences between genotypes with and without *UAS-dnTCF* (black asterisks, p<0.05, red asterisks, p<0.001) and (B) significant differences from control values (black asterisks, p<0.05, red asterisks, p<0.001).

The increased FSC division induced by increased JAK-STAT pathway activity (43.9% EdU index) was not significantly altered by addition of dnTCF (46.3%) and was only slightly reduced by loss of *arr* function (37.7%) (Fig. 5B; Fig. S3A), resembling the minimal effects of *arr* and dnTCF in a wild-type background. Ectopic division in the EC region was also largely unaltered by reducing Wnt pathway activity (16.7% vs 12.3%) (Fig. 5B). The division rate of FSCs with increased JAK-STAT pathway activity was diminished by elevating Wnt pathway activity (30.3% for *axn UAS-Hop*) but FSC division was still greater than controls (25%) and much closer to that of *UAS-Hop* FSCs (43.9%) than *axn/apc* mutant FSCs (7.4%) (Fig. 5B; Fig. S3B, C). These *axn UAS-Hop* FSCs were more prevalent in layer 1 (35.3%) than *axn/apc* mutant FSCs (20.6%), but still had an anterior bias compared to controls (48.8%) and FSCs with *UAS-Hop* alone (49.1%) (Fig. 6B-D). The proliferative phenotype of *UAS-Hop* was almost fully epistatic to *axn* when considering each FSC layer separately (35.0%, 28.6%, 24.3% in layers 1-3) (Fig. 5B).

**Figure 6.**
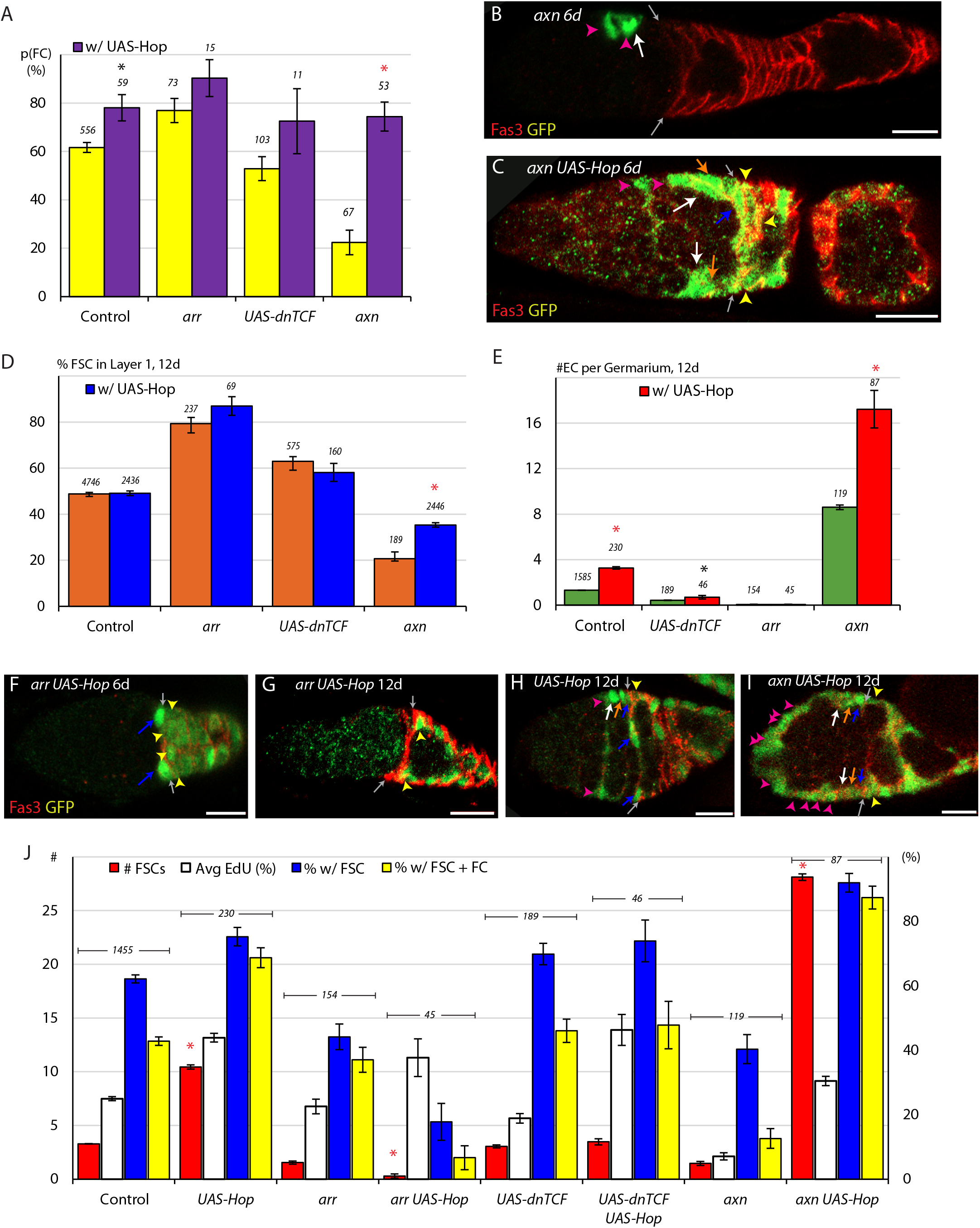
Promotion of FC production by the JAK-STAT pathway overrides the opposing influence of increased Wnt pathway activity and synergizes with loss of Wnt pathway activity to cause dramatic loss of highly proliferative FSCs. (A) Average probability of a layer 1 FSC becoming an FC during a single budding cycle for the indicated MARCM lineage genotypes with the number of informative germaria scored and significant differences resulting from the presence (purple) of UAS-Hop (black asterisks, p<0.05, red asterisks, p<0.001). (B, C) *axn* mutant MARCM lineages (green) at 6d, with the Fas3 (red) anterior border (gray arrows indicated) generally include, as here, anterior FSCs (layer 3, white arrows), and ECs (magenta arrowheads) but (C) addition of *UAS-Hop* resulted in more marked FSCs, including layer 1 (blue arrows) and layer 2 (orange arrows) FSCs, and marked FCs (yellow arrowheads). (D) Proportion of FSCs in layer 1 for the indicated MARCM lineage genotypes, with the number of FSCs scored and significant differences resulting from the presence (blue) of *UAS-Hop* (red asterisks, p<0.001). (E) Number of ECs per germarium for the indicated MARCM lineage genotypes, with the number of germaria scored and significant differences resulting from the presence (red) of *UAS-Hop* (black asterisks, p<0.05, red asterisks, p<0.001). (F-I) MARCM lineages (green) with the Fas3 (red) anterior border indicated (gray arrows) commonly showed, as here, (F) only layer 1 FSCs (blue arrows) and FCs (immediate FCs, yellow arrowheads) for *arr UAS-Hop* at 6d and (G) loss of FSCs, leaving only labeled FCs (yellow arrowheads) by 12d and (H, I) a large increase of labeled cells in EC locations (magenta arrowheads) when Wnt pathway activity is increased (*axn*) on top of increased JAK-STAT pathway (*UAS-Hop*), supplementing the many labeled FSCs in layers 1 (blue arrows), 2 (orange arrows) and 3 (white arrows) and FCs (yellow arrowheads). (J) Number of FSCs per germarium (red), frequency of FSCs incorporating EdU (aggregating all layers, white), frequency of ovarioles with a marked FSC (blue) and frequency of ovarioles with a marked FSC and marked FCs (yellow) for the indicated genotypes, with the number of germaria scored at 12d (EdU was scored at 6d) and significant differences for the number of FSCs (red asterisks, p<0.001) compared between genotypes with and without *UAS-Hop*. All scale bars are 10μm.

These results indicate that normal Wnt pathway activity inhibits FSC division under artificial conditions of removing STAT activity, even in locations (layer 1) where Wnt pathway activity is quite low. Under otherwise normal conditions, genetically increasing Wnt pathway activity to a level that approximates or slightly exceeds the highest physiological levels observed in the germarium, strongly inhibited FSC division but this inhibition was largely offset by genetically increasing JAK-STAT pathway activity. Thus, inhibitory actions of the Wnt pathway can be strong and dose-dependent but are almost entirely suppressed by JAK-STAT pathway activity under conditions when both pathways are unaltered or both are artificially elevated.

In all samples where FSCs lacked STAT activity and expressed dnTCF there was a robust posterior to anterior gradient of FSC division (Fig. 5A). This included genotypes with overall division rates close to controls (*kibra UAS-CycE stat* plus *UAS-dnTCF*; 21.6% vs 25% control EdU index). Thus, in the complete absence of one major graded signaling pathway and the near absence of another (Wnt), there remains a roughly normal FSC proliferation gradient.

### Potent acceleration of FC production from increased JAK-STAT together with absent Wnt pathway activity

Loss of STAT activity severely reduced the probability of FC production from layer 1 FSCs (from 61.6% to 36.0%), whereas loss of Wnt signaling had the converse effect, and reduction of Wnt signaling with dnTCF was without significant consequence. Reducing Wnt pathway activity with dnTCF in *stat* mutant FSCs with *kibra, wts or UAS-CycE*, slightly decreased FC production probability from 21.2% to 16.2% (Fig. S4A). Loss of both *arr* and *stat* activity cannot be tested readily.

Increased JAK-STAT activity favored FC production (78.1% probability). This was not greatly altered by the addition of dnTCF (72.5%) but was increased by loss of *arr* activity (90.3% for *arr UAS-Hop* FSCs) to a level higher than induced by Wnt signaling deficiency alone (78.9% for *arr* FSCs) (Fig. 6A). Thus, the combination of eliminating the normally low levels of Wnt signaling and increasing the already high levels of JAK-STAT pathway in layer 1 FSCs potently accelerated and almost mandated conversion to FCs. Moreover, despite accelerated loss from layer 1, 87.0% of *arr UAS-Hop* FSCs were found in this layer, representing an enhancement of the bias seen for *arr* FSCs (77.6%) (Fig. 6D), and further accelerating overall FC production. Artificially increasing Wnt pathway activity in FSCs strongly inhibited conversion to FCs (21.5% probability), but this inhibition was entirely overridden by increasing JAK-STAT pathway activity (74.4% conversion probability for *axn UAS-Hop* FSCs) (Fig. 6A). Thus, reducing Wnt pathway activity to zero synergized with elevated JAK-STAT pathway activity to potently drive FSCs to become FCs, while large increases and smaller decreases in Wnt pathway activity were without effect in the presence of elevated JAK-STAT pathway activity.

EC production was reduced for *arr* FSCs and increased dramatically for *axn* FSCs and less prominently for *stat* FSCs. Reduction of Wnt signaling with dnTCF did not reduce EC production alone or the accelerated EC production of FSCs lacking STAT activity (0.88 vs 0.87 ECs per anterior FSC from 0-6d) (Fig. S4E). The effects of *UAS-Hop* on EC production cannot readily be quantified because some of these marked cells in the EC region divide. Nevertheless, the average number of marked ECs per germarium at 12d, which was 1.3 for controls and 3.3 for *UAS-Hop*, was greatly altered for *arr UAS-Hop* (0.06) and *axn UAS-Hop* (17.2), and was slightly reduced for *UAS-dnTCF UAS-Hop* FSCs (Fig. 6E, G-I), showing that EC production is still highly responsive to changes in Wnt pathway activity in both directions even when JAK-STAT pathway activity is elevated. Thus, epistasis tests reveal a primary role for Wnt signaling in anterior regions and a primary role for JAK-STAT signaling in posterior regions.

### Compound pathway perturbations reveal potent effects of both FSC division rate and flux toward FCs on FSC competition

The relatively high number and persistence of FSCs lacking STAT activity despite low division rates has been noted earlier and attributed to diminished conversion to FCs. The addition of dnTCF increased division rates without increasing FC production from layer 1 FSCs or EC production from anterior FSCs, and might therefore be expected to increase FSC persistence. However, for *stat* alone or together with *kibra*, *wts* or *UAS-CycE* the average number of marked FSCs at 12d was not greatly altered (0.48 vs 0.33 for stat alone, 2.0 vs 2.1 for others) (Fig. S4F). The location of FSCs did shift towards layer 1 (23.6% vs 3.5% for *stat* alone, 36.9% vs 29.7% for others) (Fig. S4B-D), perhaps contributing to increased overall FC production, which was apparent from a small increase in the proportion of ovarioles with marked FSCs and FCs (14.3% vs 13.8% for stat alone, 19.0% vs 9.3% for others) (Fig. S4F). Thus, an increase in *stat* mutant FSC numbers, which might have been expected from increased division in response to dnTCF, was not observed, perhaps because increased division was offset by a modest increase in FC production. FSCs with *kibra stat* and *UAS-CycE* were already present at saturating normal numbers (15.9) by 12d, with a normal AP distribution of FCs, and that number was not significantly altered by the presence of *UAS-dnTCF* (16.9) (Fig. S4F).

Elevated JAK-STAT activity increased FSC number to 10.4 per germarium, presumably largely by increasing division rates (Fig. 6J). Division rates were only slightly reduced for *arr UAS-Hop* FSCs and unchanged for *UAS-dnTCF UAS-Hop* FSCs. However, complete Wnt pathway inhibition increased FSC depletion from anterior layers (Fig.6D) and conversion from layer 1 to FCs (Fig. 6A, F), with the net effect of drastically reducing the 12d average FSC population to 0.3 per germarium, with only 17.8% of ovarioles retaining any marked FSCs (Fig. 6G, J). Wnt pathway reduction with dnTCF only modestly increased posterior accumulation of FSCs without accelerating FC production from posterior FSCs and resulted in 3.2 FSCs per germarium (Fig. 6J). Thus, even greatly elevated FSC division cannot maintain a sufficient supply of marked FSCs when they are drained by posterior passage and conversion to FCs at the high rates promoted by high JAK-STAT and absent Wnt pathway activity.

With elevated Wnt signaling in *UAS-Hop* mutant lineages, FSC division rates remained abnormally high (*UAS-Hop* was epistatic to *axn*) (Fig. 5B), FC production from layer 1 FSCs was roughly normal (Fig. 6A) but FSCs were predominantly in anterior locations Fig. 6D), thereby reducing the overall rate of FC production relative to *UAS-Hop* alone. Accordingly, labeled FSCs accumulated to a greater extent than for *UAS-Hop* alone and indeed exceeded the normal capacity of the germarium at 28.1 *axn UAS-Hop* FSCs per germarium (Fig. 6H-J). Thus, changes in Wnt pathway activity profoundly altered the competitive success of FSCs with elevated JAK-STAT pathway activity by either promoting (loss of Wnt) or reducing (elevated Wnt) FC production, without significantly altering FSC division rates in either case.

## Discussion

Our investigations have generated a detailed picture of how a variety of fundamental FSC behaviors respond cell autonomously to positional cues relayed by the activity levels of two major signaling pathways that are graded with complementary polarities across the FSC domain. The results reveal JAK-STAT pathway activity as the primary dose-dependent agent dictating the pattern of FSC proliferation and as a major influence promoting conversion of FSCs to FCs. Wnt pathway magnitude relative to neighboring FSCs was previously shown to be a major determinant of the AP position of an FSC and its conversion to an EC (Reilein et al., 2017). Here we found that elevated Wnt pathway activity also inhibits FC production while elimination of Wnt pathway activity promotes conversion of posterior FSCs to FCs. These and other results suggest an outline for how external signals specify a functional stem cell domain and some elements of the logic for coordinating division and differentiation in a fluid collection of instantaneously heterogeneous stem cells maintained by population asymmetry.

The original perception of just two rigidly held FSCs per germarium, repeatedly undergoing asymmetric divisions to produce FCs (Margolis and Spradling, 1995) was replaced by a very different picture of 14-16 mobile FSCs of varied lifetimes, division rates and two alternative differentiation products (FCs and ECs) on the basis of detailed examination of FSC lineages over a variety of time periods (Reilein et al., 2017). The original model provided no basis for explaining why FSC maintenance was dependent on FSC division rate, why both increased and decreased Wnt signaling led to stem cell loss, or why FSC maintenance relied on input from almost every major signaling pathway tested. The revised model, in which FSC division and differentiation are independent explains why the division rate of one stem cell relative to others is a key determinant of FSC competition (Reilein et al., 2018). Likewise, we now understand that FSC loss is driven by excessive conversion of FSCs to ECs in response to elevated Wnt pathway activity and by excessive conversion of FSCs to FCs in response to loss of Wnt pathway activity (Reilein et al., 2017). We are also beginning to understand that several signals are employed to orchestrate FSC behavior because that behavior has many independent facets. Moreover, in this study, we found that the overall maintenance and amplification of an FSC can be rationalized as the sum of the individual behavioral response we measured in response to changes in major signaling pathway activities, affirming the validity of our conception of FSC behavior.

### FSC proliferation

The near-parallel gradients of FSC division frequency and JAK-STAT pathway activity, declining from posterior to anterior across the FSC domain suggested a potential causal link (Fig. 7). Indeed, when the JAK-STAT pathway was globally manipulated to be uniform across the EC and FSC domains, at a level marginally higher than seen normally in posterior FSCs, all FSCs were seen to divide at the same high rate and even some ECs divided. We also found that FSC division rate was cell autonomously and potently reduced by loss of STAT activity and increased by elevated pathway activity. Thus, the pattern of JAK-STAT pathway activity dictates the pattern of FSC divisions (Fig. 7). Moreover, the anterior border of dividing somatic cells, which is necessarily an anterior border for active stem cells in the germarium appears to be set by JAK-STAT pathway dropping below a critical threshold.

**Figure 7.**
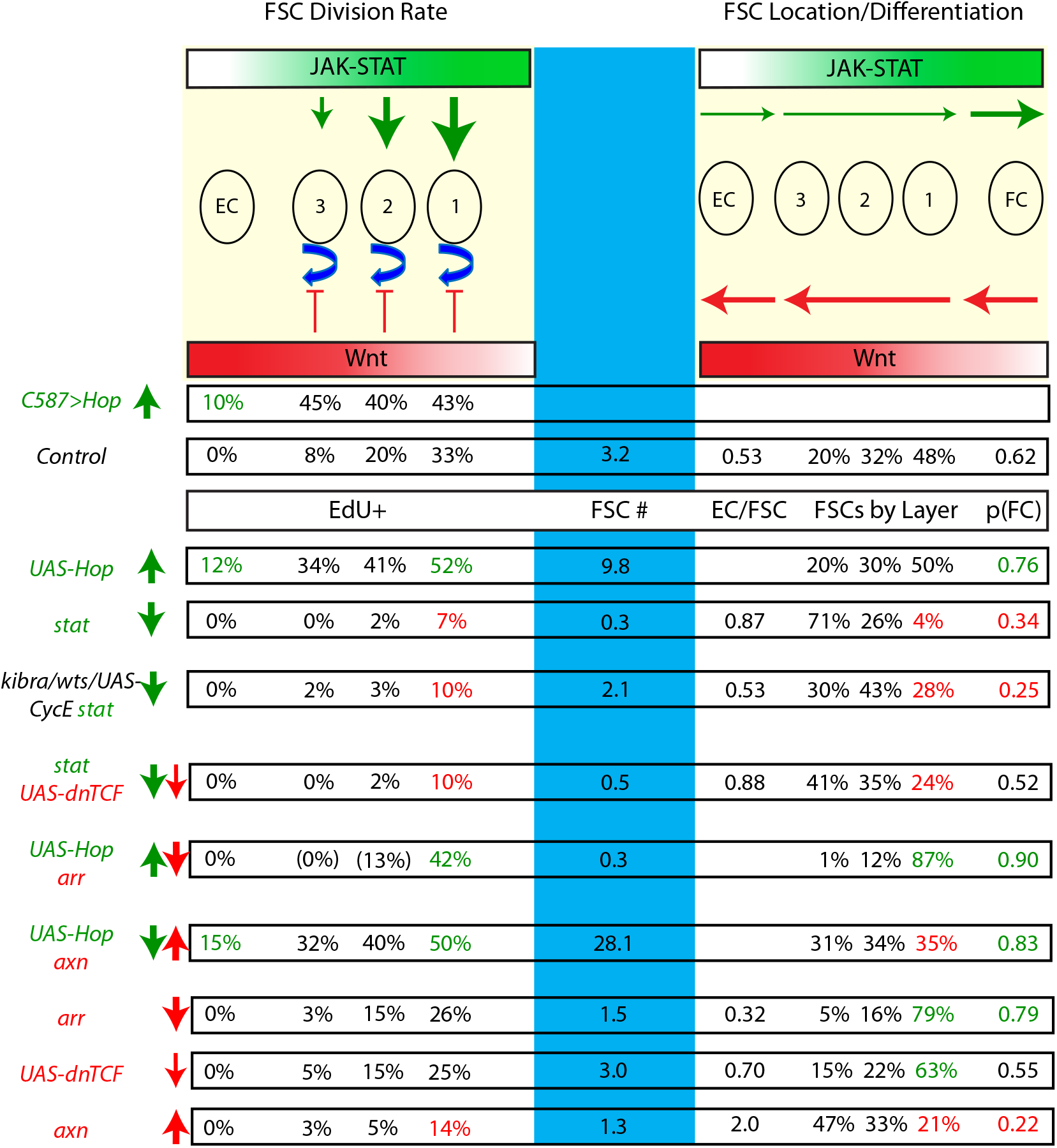
Summary of the cell autonomous influences of JAK-STAT and Wnt pathway magnitudes on FSC behavior. JAK-STAT promotes FSC division in proportion to graded pathway activity (green arrows, top), while Wnt pathway activity only inhibits division (red). FC production from posterior FSCs is promoted by JAK-STAT and loss of Wnt pathway activity. Consequently, stat mutant FSCs were maintained better than expected from their greatly reduced division rates and FSCs with high JAK-STAT activity and no Wnt pathway activity were rapidly drained despite high division rates. Increased Wnt pathway activity always favored more anterior FSC locations and EC production from anterior FSCs, and generally had a stronger influence than JAK-STAT in anterior regions. Key supporting data from global alteration of signaling (*C587*>*Hop*) and MARCM lineages are displayed with arrows indicating increases (up) or decreases (down) of JAK-STAT (green) and Wnt (red) pathway activity. Values for EdU in parentheses indicate low number of FSCs scored (n<10). Values are highlighted if significantly increased (green) or decreased (red) for EC division or FSC layer 1 behavior (EdU index, proportion of all FSCs, and FC production).

Additional influences on FSC division pattern were revealed in the absence of JAK-STAT pathway activity. FSCs lacking STAT activity always divided at a higher rate in more posterior locations when tested in five different genotype combinations, manipulating CycE and Yki pathway components to drive overall FSC division rates towards normal values. Whether the signals responsible for that polarized behavior normally augment JAK-STAT pathway action or serve only as a latent reserve system can only be tested once the signals are identified.

Graded Hh signaling promotes FSC division but it declines from anterior to posterior and cannot therefore underlie the converse gradient of FSC division (Hartman et al., 2013; Hartman et al., 2010; Huang and Kalderon, 2014; Vied et al., 2012). Genetically increased Wnt pathway activity, on the other hand, was shown previously (Reilein et al., 2017), and here, to strongly inhibit FSC division and Wnt pathway activity declines sharply from the EC domain towards FCs. Moreover, we found that in the absence of STAT activity FSC division was strongly increased when Wnt pathway activity was reduced, implying significant inhibition by normal levels of Wnt pathway activity. However, FSCs with no STAT activity and reduced Wnt pathway activity still showed a normal AP gradient of proliferation. Although Wnt pathway activity was not eliminated in these tests and residual activity may still have maintained a weak anterior to posterior gradation, it does not seem possible that Wnt pathway activity provides an essential graded proliferative influence in the absence of STAT activity.

Under conditions of normal JAK-STAT pathway activity, complete loss of Wnt pathway activity in clonal analyses mildly reduced FSC division rates and did not alter their pattern. Moreover, global reduction of Wnt pathway activity such that residual activity was likely devoid of any significant gradation, permitted a normal pattern of FSC division at roughly normal rates. The dominant influence of normal JAK-STAT pathway activity over Wnt pathway activity seen under normal conditions was observed also when both pathways were artificially elevated. These observations are not the result of cell autonomous alteration of Wnt pathway activity by JAK-STAT and will likely lead to insights about the mechanisms coupling each pathway to growth and cell cycle transitions but the physiological role of the latent influences of the Wnt pathway on FSC division (exposed under artificial genetic conditions) remain unclear. The most likely focus of influence is at the EC/FSC border where Wnt pathway activity is high and JAK-STAT pathway low. While we did not observe any ectopic EC division in response to Wnt pathway reduction or elimination, the latter observation may be limited by the possibility (discussed below) that any ECs with no Wnt pathway activity may rapidly become an FSC or die.

### FSC location and differentiation

In this study we measured FSC location and, separately, the probability that a posterior (layer 1) FSC becomes an FC. Loss of STAT activity substantially reduced the latter probability and resulted in a substantially more anterior FSC location relative to neighboring wild-type FSCs. Both factors together combine to reduce FC production. Increased JAK-STAT pathway activity increased posterior FSC to FC conversion but did not affect steady-state FSC locations. Conversely, increased Wnt pathway activity reduced FSC to FC conversion and reduced posterior FSC accumulation, while loss of Wnt pathway activity had the opposite consequences, revealing dose-dependent responses to this pathway throughout the FSC domain. Thus, through concerted actions on two separable parameters of behavior (FSC location and conversion of posterior FSCs to FCs), FC production is promoted by JAK-STAT pathway activity and reduced by Wnt pathway activity, in keeping with the relative levels of these pathways in the posterior half of the FSC domain. Although the mechanisms for FSC to FC conversion are currently unknown, it seems likely that the process is considerably more complex than the AP movements of FSCs between layers, so the concerted actions noted are not likely different manifestations of exactly the same molecular responses to signaling. The responses to loss of Wnt signaling and elevated JAK-STAT pathway activity were found to be additive, with that combination causing an extremely high rate of FC production, severely depleting the marked FSC pool. In effect, that result shows that the signaling environment posterior to layer 1 FSCs cannot support maintenance of an FSC.

EC production from anterior FSCs was measured from 0-6d after clone induction and, as noted previously, was strongly favored by increasing Wnt pathway activity. A quantitatively smaller increase was observed when STAT activity was eliminated. By measuring EC production over a longer time period (up to 12d) it became evident that even wild-type ECs produced from FSCs during adulthood had a limited lifetime, likely much shorter than that of an EC produced prior to adulthood. EC stability was further reduced by reducing or eliminating Wnt pathway activity. The co-expression of an inhibitor of apoptosis suggested that EC turnover was partly due to apoptosis but was also likely due to reversion of some ECs to FSC status, with the rates of both processes potentially increased by loss of Wnt pathway activity. The latter possibility would complement the overall picture supported by abundant evidence that all transitions in the posterior direction are favored by low Wnt pathway activity, while all anterior transitions are favored by high pathway activity (Fig. 7). It appears that loss of STAT activity universally favors anterior transitions and opposes posterior transitions but artificial increases in pathway activity appeared only to promote the FSC to FC transition (though EC production cannot be measured accurately because some cells in EC locations divide when JAK-STAT pathway activity is elevated) (Fig. 7).

### Co-ordination of FSC responses to external signals

By examining twin-spot products of recombination in an FSC (with complementary colors) where one daughter lineage consisted of just a single FC patch, it was shown that an FSC can become an FC at any time after its last division (Reilein et al., 2018). Thus, for an individual FSC there appears to be no direct causative link between division and differentiation to an FC. That separation of fundamental stem cell activities allows the possibility of independent regulation of each process at the single cell level. There may, nonetheless, be some systematic connections among individual, separable FSC behaviors.

Our results to date suggest that most behaviors are independent with the exception of an apparently modest connection between division rate and AP location. We found that *cycE* mutant FSCs, which likely have reduced division as the only direct consequence, had a significant anterior bias and that *stat* mutant FSCs showed a reduced anterior bias when division rate was increased by elevating CycE expression or Yki activity. The secondary AP displacement might plausibly be caused by poorer competition of slow-dividing FSCs in the posterior layer where normal FSCs divide and become FCs at a high rate.

In contrast to the effect of division rate on AP location, the greatly reduced FSC to FC conversion of *stat* mutants was not altered by increasing division rate, and the increased conversion of FSCs to FCs due to elevated JAK-STAT pathway activity was not altered by reducing FSC division rates (with a Cdk2 inhibitor). Hence, there must be specific mechanisms that balance FSC division and differentiation because this is not accomplished by robustly connecting these properties at the single cell level. Balance could be achieved in theory if a key signal affected both division and differentiation in appropriate proportions. An important parallel consideration is how signals define the domain in which stem cells can exist, thereby defining stem cell numbers.

The JAK-STAT pathway promotes both FSC division and conversion to FCs. About 5-6 FCs are produced per 12h budding cycle, while only one fourth as many anterior FSCs become ECs. Conversion to FCs is therefore the main drain on the FSC population that mandates compensatory FSC division. Because JAK-STAT signaling stimulates each process in a dose-dependent manner, maintenance of an individual FSC without amplification will be buffered against stochastic or systemic changes in the strength of this pathway. Also, by instructing posterior FSCs to divide faster than anterior FSCs the generation and loss of FSCs is roughly balanced for each layer, allowing for a roughly equal flux of FSCs between layers, rather than, for example, a net anterior to posterior flow that would make anterior FSCs systematically longer-lived. The contribution of JAK-STAT signaling to coordination of FSC division and differentiation was evident when only the division rate of stat mutant FSCs was elevated by genetic alteration of other agents. The consequence was a large accumulation of hyper-competitive marked FSCs that supported very little FC production.

There does not appear to be any comparable dual response co-ordination for the Wnt pathway. Higher levels than normal promote anterior FSC movement and reduce conversion to FCs, lower levels promote posterior FSC movement and conversion to FCs, with neither response inducing increased FSC division. Nevertheless, it is quite striking that loss of Wnt pathway activity alone allows FSCs to be retained over 12d in a fairly high proportion of ovarioles (44.2% vs 62.1% in controls). However, the FSC population was drained rapidly when FSCs additionally had elevated JAK-STAT pathway activity (Fig. 7). Thus, normal Wnt pathway activity plays a supporting role in allowing co-ordinated proliferative and differentiation responses to altered JAK-STAT signaling that preserve FSCs.

The division rate appropriate to support production of a specific number of FCs and ECs every 12h also depends on the total number of FSCs. This, in turn, appears to be defined by a specific domain or space where FSCs reside. Both graded JAK-STAT and Wnt pathways contribute to both borders. The anterior border is between non-dividing ECs and FSCs. Increasing Wnt pathway activity and decreased JAK-STAT pathway activity both promote FSC to EC conversion, while declining JAK-STAT signaling also limits the proliferative zone. The posterior border is between FSCs and FCs. Increased JAK-STAT and reduction of Wnt pathway activity (to zero) promote the FSC to FC transition. These results and assertions apply to cell autonomous responses examined in mosaic tissues (where most cells have normal genotypes). Further experiments manipulating pathway activities globally (in all cells) may well result in a number of compensatory changes in behavior and signaling properties, and will have to be evaluated in detail to understand to what extent the size and location of the FSC domain depends on the normal magnitude and gradations of these and other pathways.

## Acknowledgments

This work was supported by NIH RO1 GM079351 to DK. We thank Aaron Choi, Amy Reilein, Jamie Little, Jianhua Huang, Haoran Liu and Diana Kim for research assistance, Amy Reilein, Rachel Misner, Lena Kogan, Tulle Hazelrigg and Iva Greenwald for continued discussions and input, the Bloomington stock center for provision of genetic reagents, the Developmental Studies Hybridoma Bank (DSHB) for antibodies, FlyBase as an information resource, and the confocal microscope resource provided by the Dept. of Biological Sciences, Columbia University.

## Methods

### MARCM Clonal Analysis

1-3d old adult *Drosophila melanogaster* females with the appropriate genotypes were given a single 30 min (for *FRT40A* and *FRT42D*) or 45 min (for *FRT82B*) heat shock at 37C, with the heat shock duration determined by the observed relative rate of recombination per unique FRT sites. Afterwards, flies were incubated at either 25C or 29C. The two temperatures were used to compare levels of *UAS-GAL4* driven activity. Flies were maintained by frequent passage on normal rich food supplemented by fresh wet yeast during the 12d experimental period. Flies were dissected at 6d and 12d. We waited until 6d to ensure that all GAL80 present in cells, prior to clone induction, would be titrated out, permitting robust GAL4 induction of *UAS-GFP* and any additional transgenes. The 6d time point also ensured that any marked cells are derived from FSCs, as dividing FCs marked in the heat shock would have passed through the ovariole in less than 5d.

Immediately after dissection, 6d ovaries underwent 1h of EdU labelling based on the protocol of the Click-iT™ Plus EdU Cell Proliferation Kit for Imaging (Invitrogen). Both 6d and 12d ovaries were stained for Fasciclin III (Fas3) and GFP. Ovaries were then manually separated into constituent ovarioles, and mounted using DAPI Fluoromount-G® (SouthernBiotech) to stain nuclei. Ovarioles were imaged with a Zeiss LSM700 or LSM800 confocal microscope, operated in part by the Zeiss ZEN software. The entire germarium was captured in the images, as well as an average of 3-4 egg chambers. Collected images were saved as CZI files, and were later analyzed utilizing the ZEN Lite software. We aimed to image 50 germaria for every genotype in each experiment.

### MARCM Genotypes

Flies with alleles on an *FRT40A*, *FRT42D*, and *FRT82B* chromosomes were used in MARCM experiments:

FRT40A: *hs-Flp, UAS-nGFP, tub-GAL4; act-GAL80 FRT40A / (X)FRT40A); act>CD2>GAL4/UAS-(Y)* – where X,Y combinations included: (X) – *NM* (Nuclear Myc, Control), *cycE^WX^*, *cutlet^4.5.43^* (Y) - *UAS-Hop^3W^, UAS-Dap, UAS-dnTCF*
FRT42D: *hs-Flp, UAS-nGFP, tub-GAL4; FRT42D act-GAL80 tub-GAL80 / FRT42D (X); act>CD2>GAL4/ UAS-(Y)* – where X,Y combinations included: (X) – *sha* (Control), *ubi-GFP* (Control), *arr^2^,* (Y) – *UAS-Hop^3W^, UAS-DIAP1*
FRT82B: *hs-Flp, UAS-nGFP, tub-GAL4; act>CD2>GAL4 UAS-GFP / UAS-(Y); FRT82B tub-GAL80/FRT82B (X)* – where X,Y combinations included: (X) – *NM* (control), *stat^085C9^, stat^06346^, axn^E77^, axn^S044320^, apc1^Q8^apc2^D40^, kibra^32^, kibra^del^, wts^x1^,* (Y) *UAS-Hop^3W^, UAS-CycE, UAS-dnTCF*

### C587-GAL4 Experiments

1-3d old flies of the genotype *C587-GAL4; UAS-X/ts-GAL80, FRT42D tub-lacZ; (Reporter)/TM6B* were chosen, where *UAS-X* was *UAS-dnTCF*, *UAS-CycE*, or *UAS-Hop*, and the reporter was either Stat-GFP or Fz3-RFP. Flies were incubated at 29C for 3d, and *UAS-Hop* flies were also incubated for 6d and 10d. Dissected ovaries underwent the EdU and Immunohistochemistry protocols as above, without staining for GFP. For Fz3-RFP experiments with EdU, Alexa Fluor™ 488 dye was used instead of 594 to avoid spectral overlap.

### EdU Protocol

Ovaries were directly dissected into a solution of 15 μM EdU in Schneider’s *Drosophila* media (500μl, Gibco) for one hour at room temperature. These tubes were laid on their side and rocked manually, to ensure all dissected ovaries were fully submerged. Ovaries were then fixed in 3.7% paraformaldehyde in PBS for 10 minutes, treated with Triton in PBS (500 μl, 0.5% v/v) for 20 minutes, and rinsed 2x with bovine serum albumin (BSA) in PBS (500 μl, 3% w/v) for 5 minutes each rinse. Ovaries were exposed to the Click-iT Plus reaction cocktail (500 μl) for EdU visualization, for 45 minutes. The reaction cocktail was freshly prepared prior to use, with reagents from the Invitrogen™ Click-iT™ Plus EdU Cell Proliferation Kit for Imaging, including the Alexa Fluor™ 594 dye. Ovaries were then rinsed 3x with BSA in PBS (500 μl, 3% w/v) for 5 minutes each rinse.

### Immunohistochemistry

For experiments without EdU, ovaries were dissected directly into a fixation solution of 4% paraformaldehyde in PBS for 10 min at room temperature, rinsed 3x in PBS, and blocked in 10% normal goat serum (NGS) (Jackson ImmunoResearch Laboratories) in PBS with 0.1% Triton and 0.05% Tween-20 (PBST) for 1 h. Monoclonal antibodies for Fas-3 were obtained from the Developmental Studies Hybridoma Bank, created by the NICHD of the NIH and maintained at The University of Iowa, Department of Biology, Iowa City, IA 52242. 7G10 anti-Fasciclin III was deposited to the DSHB by Goodman, C. and was used at 1:250. Other primary antibodies used were anti-GFP (A6455, Molecular Probes) at 1:1000. Ovaries were incubated in primary antibodies overnight, rinsed three times in PBST, and incubated 1-2 h in secondary antibodies Alexa-488 and Alexa-647 (ThermoFisher) to label GFP and Fas3, respectively. DAPI-Fluoromount-G (Southern Biotech) was used to mount ovaries.

### Imaging and scoring

All germaria were imaged in three dimensions on an LSM700 or LSM800 confocal laser scanning microscope (Zeiss) and using a 63x 1.4 N.A. lens. Zeiss ZEN software was used to operate the microscope and view images. Images were typically 700×700 pixels with a bit depth of 12. The scaling per pixel was 0.21 μm x 0.21 μm x 2.5 μm. The range indicator in ZEN was used to determine the appropriate laser intensity and gain. ZEN was used to linearly adjust channel intensity for dim signals to improve brightness without photobleaching samples. Images were saved as CZI files and scored directly in ZEN. DAPI and Fas3 staining were used as landmarks to guide scoring. Marked cells were considered FSCs if they were within three cell diameters anterior of the Fas3 border. Cells immediately adjacent to the border were considered to be in Layer 1, with Layers 2 and 3 in sequentially anterior positions. Anterior to the FSC niche, the EC region was roughly divided into two halves, with region 2a ECs immediately anterior to FSCs and region 1 ECs anterior to that. Germaria were also scored (Y/N) for the presence of marked FCs. For the “Immediate FC Method” (Appendix) tabulation, the presence of an FC immediately posterior to Layer 1 was also scored Y/N. For publication, images were digitally zoomed in ZEN and exported as tif files using the “Contents of Image Window” function. Images were rotated in Abode Photoshop CS5 to uniformly orient the germaria.

### Measurement of signaling pathway reporter activities

Stat-GFP and Fz3-RFP reporter activity was quantified within ZEN software. Using the Draw Spline Contour function, an outline of a DAPI cell nuclei was traced, and the fluorescence intensity within the outline was recorded. The outline of cells not expressing the reporter were used to determine background intensity and were subtracted from calculated totals. For quantification of signaling pathway gradients, the intensity of FCs from the second or third egg chamber was used as a reference, and all intensity measurements of cells within the germarium were divided by the reference to produce a relative intensity that could be compared. For quantification of individual clones, the intensity of a GFP-positive cell was divided by that of a GFP-negative cell in a similar position along the A/P axis, within the same or an adjacent Z plane and an average was calculated from many such pairs to derive the percentage intensity for labeled cells relative to unlabeled cells.

### Statistics and Reproducibility

All images shown are representative of at least ten examples. In most cases the number is much higher and is given explicitly where relevant for statistical analysis of outcomes. No statistical method was used to predetermine sample size but we used prior experience to establish minimal sample sizes. No samples were excluded from analysis, provided staining was of high quality. The experiments were not randomized; all samples presented as groups in the results were part of the same experiment and treated in exactly analogous ways without regard to the identity of the sample. Investigators were not blinded during outcome assessment, but had no pre-conception of what the outcomes might be. For EdU incorporation, FSC layers, Immediate FC probability tabulations, proportion of germaria with FSCs and/or FCs, the “N-1” Chi-squared test method was used to calculate a Z score for determining significance between indicated genotypes, and error was reported as standard error of a proportion. For average number of FSCs, Fz3-RFP reporter intensity comparison, and EC/FSC ratio, a t-test was used to determine significance between indicated genotypes, and error was reported as standard error of the mean.

### Data Availability

All data supporting the findings from this study are available from the corresponding author upon reasonable request.

### Immediate FC Method for calculating posterior FSC to FC conversion probability

The immediate FC method was used to calculate the probability for any layer 1 FSC to become an FC in a given cycle of egg chamber budding. As layer 1 FSCs directly give rise to FC daughters (Reilein et al, 2017), this was assessed by determining the proportion of germaria that contained a marked FC immediately posterior to the FSC region, which indicated recent FC production. This was only assessed in germaria with a small number of FSCs (1-3) to reasonably deduce the likelihood for an individual FSC. As the rate of proliferation would also influence this probability, this was accounted for in the immediate FC method equation.

Probability of a single layer 1 FSC becoming an FC in one cycle = p
Probability of a single layer 1 FSC dividing in one cycle = q

We assume that on average FSC division occurs halfway through a cycle, such that the probability of a newly-produced FSC becoming an FC is p/2. Therefore, the total probability (P) of an FSC becoming an FC in one cycle is the sum of two probabilities: an FSC becoming an FC and an FSC dividing and the additional FSC becoming an FC.

P = p + (1-p)*q*(p/2)
P = p + pq/2 – p^2^q/2

We calculate the probability of an FSC becoming an FC by tallying the proportion of germaria (x) that do not have immediate FC daughters when a single marked layer 1 FSC is present (we assume that, on average, a single marked layer 1 FSC was present at the start of the prior cycle).

x = 1 – (p + pq/2 – p^2^q/2)
x = 1 – p – pq/2 + p^2^q/2
x = 1 – (1+q/2)*p + (q/2)*p^2^
0 = (q/2)*p^2^ – (1+q/2)*p + (1-x)
p = [(1+q/2) +/− SQRT((-(1+q/2)^2^) – (4(q/2)(1-x)))]/q (using quadratic formula)

Using this equation, we can solve for p, as x and q can be tabulated from scoring data. We also considered germaria with immediate FC daughters that have no FSCs present in layer 1, as the only possibility is that a layer 1 FSC was present at the start of the last cycle and then became an FC. These instances were incorporated into the calculation of the proportion of germaria with a single layer 1 FSC but no immediate FCs.

If 2 or 3 FSCs were present, the square or cube root of the x ratio was used, respectively. A weighted average of adjusted x ratios for germaria with 1, 2 and 3 FSCs was calculated (weighted according to the number of examples of 0-1, 2 and 3 layer 1 FSCs) and used in the formula to calculate p.

The q value was adjusted based on measured proliferation for mutants compared to controls, as well as predicted daughter cell production, which assumes that seven FSCs (out of a total of 16 FSCs) are dividing per cycle to produce 5.6 FCs and 1.4 ECs per cycle. Therefore: q = “Proportion of EdU incorporation for layer 1 FSCs (mutant or control)”/”average EdU incorporation of all control FSCs” * 7/16.

### EC per anterior FSC ratio

The EC/anterior FSC ratio was calculated by dividing the total number of marked ECs by the total number of marked anterior FSCs in all germaria that contained at least one marked FSC (in any position), at both 6d and 12d. For mutant genotypes, an average of the control anterior FSC number and the mutant anterior FSC number was used for the anterior FSC number, approximating the average number of marked mutant anterior FSCs during the experimental time period (because control FSC numbers should be constant throughout on average, and should be the same as mutant FSC numbers just after clone induction). For controls, an unweighted average of all ratios from 31 experiments was reported as the final EC production ratio values at 6d and 12d. There was some variability in the measurements, so we chose to use an unweighted average to prevent any particular experiment to be overly represented in the final value, since there was a similar number of germaria scored in each replicate. For mutant genotypes, a weighted average was reported for samples with multiple replicates. We also calculated the % change between 6d and 12d values with the formula (12d-6d)/6d.

## References

Bach, E.A., Ekas, L.A., Ayala-Camargo, A., Flaherty, M.S., Lee, H., Perrimon, N., and Baeg, G.H. (2007). GFP reporters detect the activation of the Drosophila JAK/STAT pathway in vivo. Gene Expr Patterns 7, 323–331.

Castanieto, A., Johnston, M.J., and Nystul, T.G. (2014). EGFR signaling promotes self-renewal through the establishment of cell polarity in Drosophila follicle stem cells. Elife 3.

Clevers, H., and Watt, F.M. (2018). Defining Adult Stem Cells by Function, not by Phenotype. Annu Rev Biochem 87, 1015–1027.

Decotto, E., and Spradling, A.C. (2005). The Drosophila ovarian and testis stem cell niches: similar somatic stem cells and signals. Developmental Cell 9, 501–510.

Duhart, J.C., Parsons, T.T., and Raftery, L.A. (2017). The repertoire of epithelial morphogenesis on display: Progressive elaboration of Drosophila egg structure. Mech Dev 148, 18–39.

Forbes, A.J., Spradling, A.C., Ingham, P.W., and Lin, H. (1996). The role of segment polarity genes during early oogenesis in Drosophila. Development 122, 3283–3294.

Fox, D.T., Morris, L.X., Nystul, T., and Spradling, A.C. (2008). Lineage analysis of stem cells. In StemBook (Cambridge (MA)).

Greulich, P., and Simons, B.D. (2016). Dynamic heterogeneity as a strategy of stem cell self-renewal. Proc Natl Acad Sci U S A 113, 7509–7514.

Hartman, T.R., Strochlic, T.I., Ji, Y., Zinshteyn, D., and O’Reilly, A.M. (2013). Diet controls Drosophila follicle stem cell proliferation via Hedgehog sequestration and release. J Cell Biol 201, 741–757.

Hartman, T.R., Zinshteyn, D., Schofield, H.K., Nicolas, E., Okada, A., and O’Reilly, A.M. (2010). Drosophila Boi limits Hedgehog levels to suppress follicle stem cell proliferation. J Cell Biol 191, 943–952.

Hayashi, Y., Yoshinari, Y., Kobayashi, S., and Niwa, R. (2020). The regulation of Drosophila ovarian stem cell niches by signaling crosstalk. Curr Opin Insect Sci 37, 23–29.

Huang, J., and Kalderon, D. (2014). Coupling of Hedgehog and Hippo pathways promotes stem cell maintenance by stimulating proliferation. J Cell Biol 205, 325–338.

Johnston, M.J., Bar-Cohen, S., Paroush, Z., and Nystul, T.G. (2016). Phosphorylated Groucho delays differentiation in the follicle stem cell lineage by providing a molecular memory of EGFR signaling in the niche. Development 143, 4631–4642.

Jones, P.H. (2010). Stem cell fate in proliferating tissues: equal odds in a game of chance. Dev Cell 19, 489–490.

Kirilly, D., Spana, E.P., Perrimon, N., Padgett, R.W., and Xie, T. (2005). BMP signaling is required for controlling somatic stem cell self-renewal in the Drosophila ovary. Developmental Cell 9, 651–662.

Kirilly, D., Wang, S., and Xie, T. (2011). Self-maintained escort cells form a germline stem cell differentiation niche. Development 138, 5087–5097.

Kretzschmar, K., and Watt, F.M. (2012). Lineage tracing. Cell 148, 33–45.

Lane, M.E., Sauer, K., Wallace, K., Jan, Y.N., Lehner, C.F., and Vaessin, H. (1996). Dacapo, a cyclin-dependent kinase inhibitor, stops cell proliferation during Drosophila development. Cell 87, 1225–1235.

Lee, T., and Luo, L. (2001). Mosaic analysis with a repressible cell marker (MARCM) for Drosophila neural development.[erratum appears in Trends Neurosci 2001 Jul;24(7):385]. Trends in Neurosciences 24, 251–254.

Lehner, C.F., Ried, G., Stern, B., and Knoblich, J.A. (1992). Cyclins and cdc2 kinases in Drosophila: genetic analyses in a higher eukaryote. Ciba Found Symp 170, 97–109; discussion 110-104.

Luo, L., Wang, H., Fan, C., Liu, S., and Cai, Y. (2015). Wnt ligands regulate Tkv expression to constrain Dpp activity in the Drosophila ovarian stem cell niche. J Cell Biol 209, 595–608.

Margolis, J., and Spradling, A. (1995). Identification and behavior of epithelial stem cells in the Drosophila ovary. Development 121, 3797–3807.

McGregor, J.R., Xi, R., and Harrison, D.A. (2002). JAK signaling is somatically required for follicle cell differentiation in Drosophila. Development 129, 705–717.

Mesa, K.R., Kawaguchi, K., Cockburn, K., Gonzalez, D., Boucher, J., Xin, T., Klein, A.M., and Greco, V. (2018). Homeostatic Epidermal Stem Cell Self-Renewal Is Driven by Local Differentiation. Cell Stem Cell 23, 677–686 e674.

O’Reilly, A.M., Lee, H.H., and Simon, M.A. (2008). Integrins control the positioning and proliferation of follicle stem cells in the Drosophila ovary. J Cell Biol 182, 801–815.

Post, Y., and Clevers, H. (2019). Defining Adult Stem Cell Function at Its Simplest: The Ability to Replace Lost Cells through Mitosis. Cell Stem Cell 25, 174–183.

Pritchett, T.L., Tanner, E.A., and McCall, K. (2009). Cracking open cell death in the Drosophila ovary. Apoptosis 14, 969–979.

Reilein, A., Melamed, D., Park, K.S., Berg, A., Cimetta, E., Tandon, N., Vunjak-Novakovic, G., Finkelstein, S., and Kalderon, D. (2017). Alternative direct stem cell derivatives defined by stem cell location and graded Wnt signalling. Nat Cell Biol 19, 433–444.

Reilein, A., Melamed, D., Tavare, S., and Kalderon, D. (2018). Division-independent differentiation mandates proliferative competition among stem cells. Proc Natl Acad Sci U S A 115, E3182–E3191.

Ritsma, L., Ellenbroek, S.I., Zomer, A., Snippert, H.J., de Sauvage, F.J., Simons, B.D., Clevers, H., and van Rheenen, J. (2014). Intestinal crypt homeostasis revealed at single-stem-cell level by in vivo live imaging. Nature 507, 362–365.

Rompolas, P., Mesa, K.R., Kawaguchi, K., Park, S., Gonzalez, D., Brown, S., Boucher, J., Klein, A.M., and Greco, V. (2016). Spatiotemporal coordination of stem cell commitment during epidermal homeostasis. Science 352, 1471–1474.

Sahai-Hernandez, P., and Nystul, T.G. (2013). A dynamic population of stromal cells contributes to the follicle stem cell niche in the Drosophila ovary. Development 140, 4490–4498.

Simons, B.D., and Clevers, H. (2011). Strategies for homeostatic stem cell self-renewal in adult tissues. Cell 145, 851–862.

Snippert, H.J., van der Flier, L.G., Sato, T., van Es, J.H., van den Born, M., Kroon-Veenboer, C., Barker, N., Klein, A.M., van Rheenen, J., Simons, B.D., et al. (2010). Intestinal crypt homeostasis results from neutral competition between symmetrically dividing Lgr5 stem cells. Cell 143, 134–144.

Song, X., Wong, M.D., Kawase, E., Xi, R., Ding, B.C., McCarthy, J.J., and Xie, T. (2004). Bmp signals from niche cells directly repress transcription of a differentiation-promoting gene, bag of marbles, in germline stem cells in the Drosophila ovary. Development 131, 1353–1364.

Song, X., and Xie, T. (2003). Wingless signaling regulates the maintenance of ovarian somatic stem cells in Drosophila. Development 130, 3259–3268.

Spencer, S.L., Cappell, S.D., Tsai, F.C., Overton, K.W., Wang, C.L., and Meyer, T. (2013). The proliferation-quiescence decision is controlled by a bifurcation in CDK2 activity at mitotic exit. Cell 155, 369–383.

van de Wetering, M., Sancho, E., Verweij, C., de Lau, W., Oving, I., Hurlstone, A., van der Horn, K., Batlle, E., Coudreuse, D., Haramis, A.P., et al. (2002). The beta-catenin/TCF-4 complex imposes a crypt progenitor phenotype on colorectal cancer cells. Cell 111, 241–250.

Vied, C., Reilein, A., Field, N.S., and Kalderon, D. (2012). Regulation of stem cells by intersecting gradients of long-range niche signals. Dev Cell 23, 836–848.

Waghmare, I., and Page-McCaw, A. (2018). Wnt Signaling in Stem Cell Maintenance and Differentiation in the Drosophila Germarium. Genes (Basel) 9.

Wang, S., Gao, Y., Song, X., Ma, X., Zhu, X., Mao, Y., Yang, Z., Ni, J., Li, H., Malanowski, K.E., et al. (2015). Wnt signaling-mediated redox regulation maintains the germ line stem cell differentiation niche. Elife 4, e08174.

Wang, X., and Page-McCaw, A. (2014). A matrix metalloproteinase mediates long-distance attenuation of stem cell proliferation. J Cell Biol 206, 923–936.

Wang, X., and Page-McCaw, A. (2018). Wnt6 maintains anterior escort cells as an integral component of the germline stem cell niche. Development 145.

Wang, Z.A., Huang, J., and Kalderon, D. (2012). Drosophila follicle stem cells are regulated by proliferation and niche adhesion as well as mitochondria and ROS. Nature communications 3, 769.

Wang, Z.A., and Kalderon, D. (2009). Cyclin E-dependent protein kinase activity regulates niche retention of Drosophila ovarian follicle stem cells. Proc Natl Acad Sci U S A 106, 21701–21706.

Xi, R., McGregor, J.R., and Harrison, D.A. (2003). A gradient of JAK pathway activity patterns the anterior-posterior axis of the follicular epithelium. Developmental Cell 4, 167–177.

Zeidler, M.P., Tan, C., Bellaiche, Y., Cherry, S., Hader, S., Gayko, U., and Perrimon, N. (2004). Temperature-sensitive control of protein activity by conditionally splicing inteins. Nature biotechnology 22, 871–876.

Zhang, Y., and Kalderon, D. (2001). Hedgehog acts as a somatic stem cell factor in the Drosophila ovary. Nature 410, 599–604.

