## Supplemental Figures for "Opposing Wnt and JAK-STAT signaling gradients define a stem cell domain by regulating spatially patterned cell division and differentiation at two borders"

**Supplementary Figure Legends**

**Figure S1. JAK-STAT pathway promotes division of FSCs and cells in EC territory.**

(A, B) EdU (blue) incorporation into MARCM FSC lineages expressing *UAS-Hop* 6d after clone induction with anterior border of Fas3 (red) marked (gray arrows), showing (A, B) GFP and (A', B') EdU channels separately. (A) The proportion of all GFP-positive FSCs (white and yellow arrows) that incorporated EdU (yellow arrows) was higher than for controls and (B) there were sometimes GFP-positive cells in EC locations that incorporated EdU (magenta arrowhead), which are never seen for control MARCM lineages. All scale bars are 10µm.

**Figure S2: Fz3-RFP intensity is not affected by JAK-STAT activity.**

(A) Average *Fz3-RFP* Wnt pathway reporter intensity in MARCM lineages of the indicated genotypes compared to unmarked neighbors at the same AP location and in the same z-section (set at 100% and indicated by the dashed line) with the number of cells measured above the columns. A second dashed line (at 39%) represents the RFP intensity observed in *arr* mutant cells, which are expected to have no Wnt pathway activity (39%). (B) Images showing the outlines of GFP-positive cells (long dashed line) of the indicated genotypes and their unmarked neighbor (short dashed line) traced from DAPI nuclear staining (white) to evaluate Fz3-RFP (red) intensity. Fas3 (blue) and all scale bars are 10µm.

**Figure S3. Increased JAK-STAT signaling promotes FSC division even when Wnt pathway activity is increased.**

(A-C) EdU (blue) incorporation into MARCM FSC lineages 6d after clone induction with anterior border of Fas3 (red) marked (gray arrows), showing (A-C) GFP and (A'-C') EdU channels separately. The proportion of marked FSCs with EdU (yellow arrows) compared to those without EdU (white arrows) was (A, B) higher than controls for *arr* and *axn* genotypes with *UAS-Hop* and (C) lower than controls for *axn* with no alteration of JAK-STAT signaling. All scale bars are 10µm.

**Figure S4. Decreasing Wnt signaling increases FSC division and posterior location but does not strongly alter loss of FC production or increased FC production in the absence of JAK-STAT signaling.**

(A) Average probability of a layer 1 FSC becoming an FC during a single budding cycle for the indicated MARCM lineage genotypes with the number of informative germaria scored and significant differences resulting from the presence (purple) of *UAS-dnTCF* (black asterisks, $p<0.05$). (B, C) Reduction of Wnt signaling with *UAS-dnTCF* increased the very low incorporation of EdU (blue) into (B) *stat* and (C) *UAS-CycE stat* mutant FSC lineages (green). GFP-positive cells with EdU (orange arrows) and without EdU (blue arrow) are indicated together with the Fas3 (red) anterior border (gray arrows). (D) Proportion of FSCs in layer 1 for the indicated MARCM lineage genotypes, with the number of FSCs scored and significant differences resulting from the presence (blue) of *UAS-dnTCF* (red asterisks, $p<0.001$). (E) ECs produced per anterior FSC from 0-6d for the indicated MARCM lineage genotypes with the total number of informative germaria scored. (F) Number of FSCs per germarium (red), frequency of FSCs incorporating EdU (aggregating all layers, white), frequency of ovarioles with a marked FSC (blue) and frequency of ovarioles with a marked FSC and marked FCs (yellow) for the indicated genotypes, with the number of germaria scored at 12d (EdU was scored at 6d) and significant differences for the number of FSCs (black asterisks, $p<0.05$) compared between genotypes with and without *UAS-dnTCF*. All scale bars are 10μm.

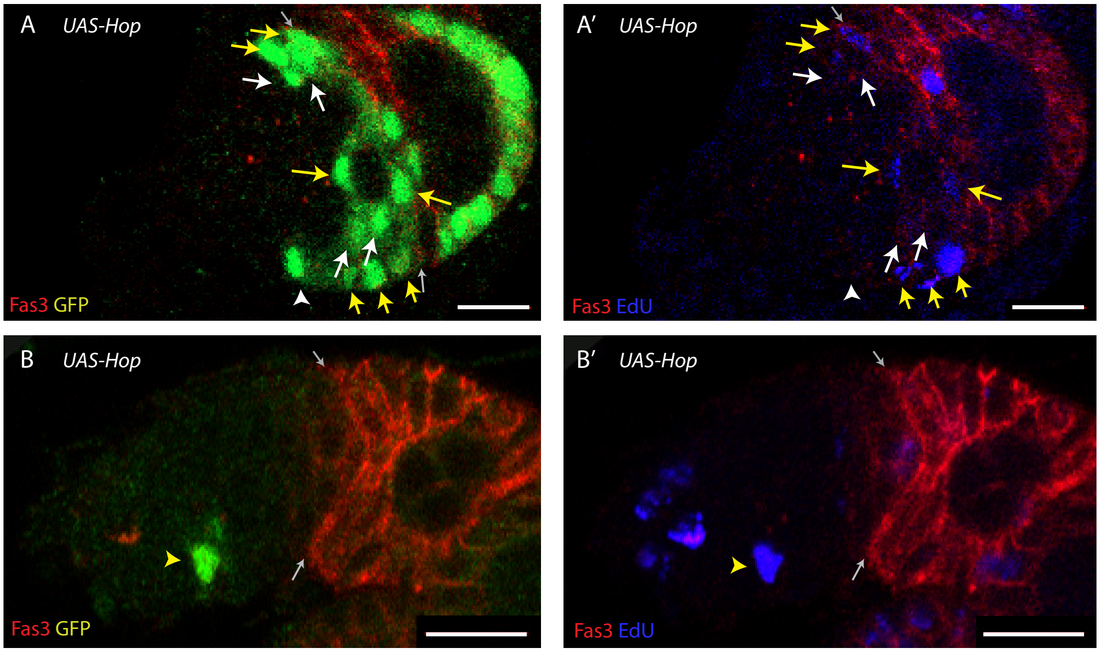
A
UAS-Hop
Fas3 GFP
A'
UAS-Hop
Fas3 EdU
B
UAS-Hop
Fas3 GFP
B'
UAS-Hop
Fas3 EdU

Figure S1

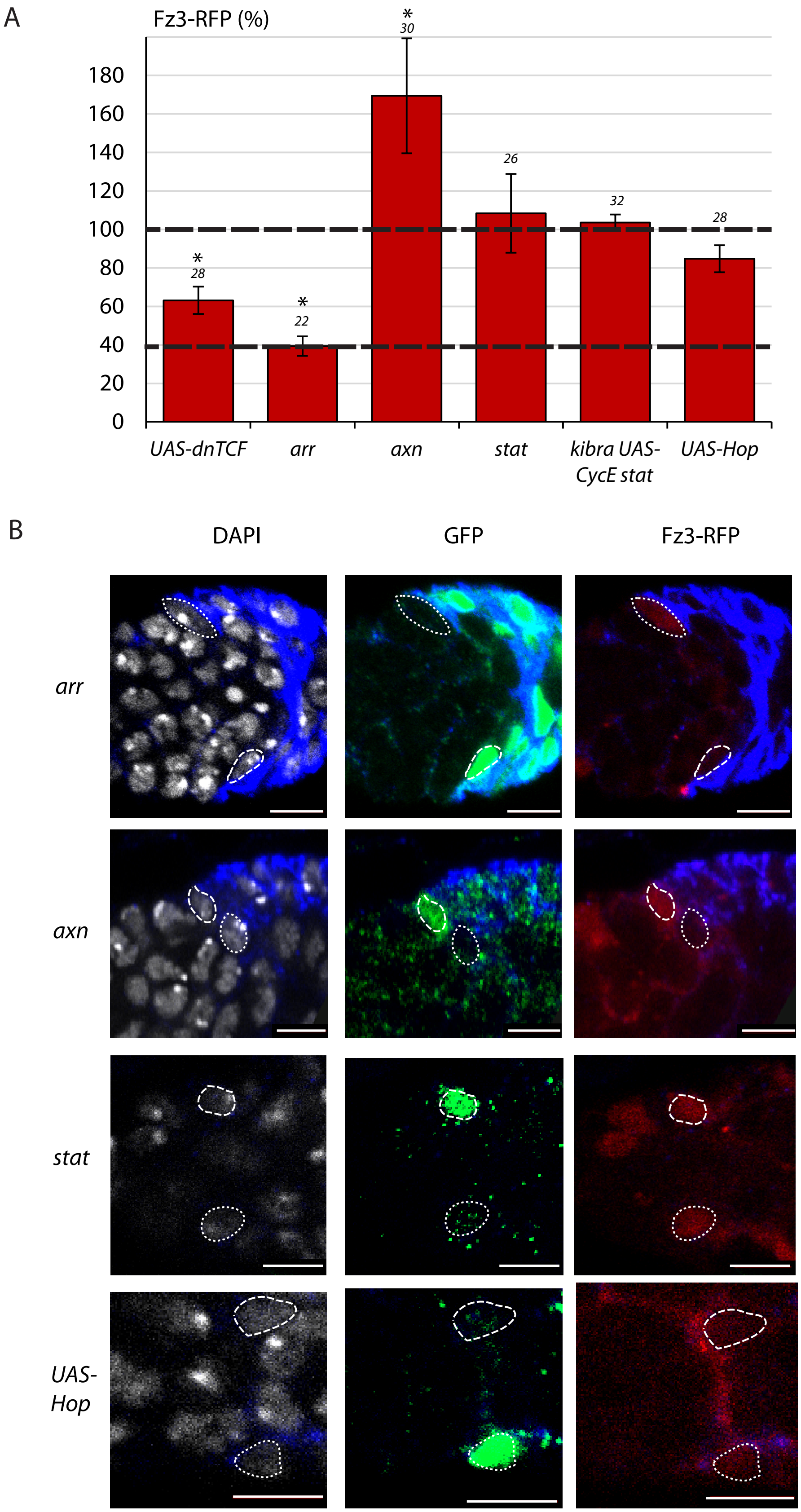
A
Fz3-RFP (%)
0
20
40
60
80
100
120
140
160
180
*
28
*
22
*
30
26
32
28
UAS-dnTCF
arr
axn
stat
kibra UAS-CycE stat
UAS-Hop
B
DAPI
GFP
Fz3-RFP
arr
axn
stat
UAS-Hop

Figure S2

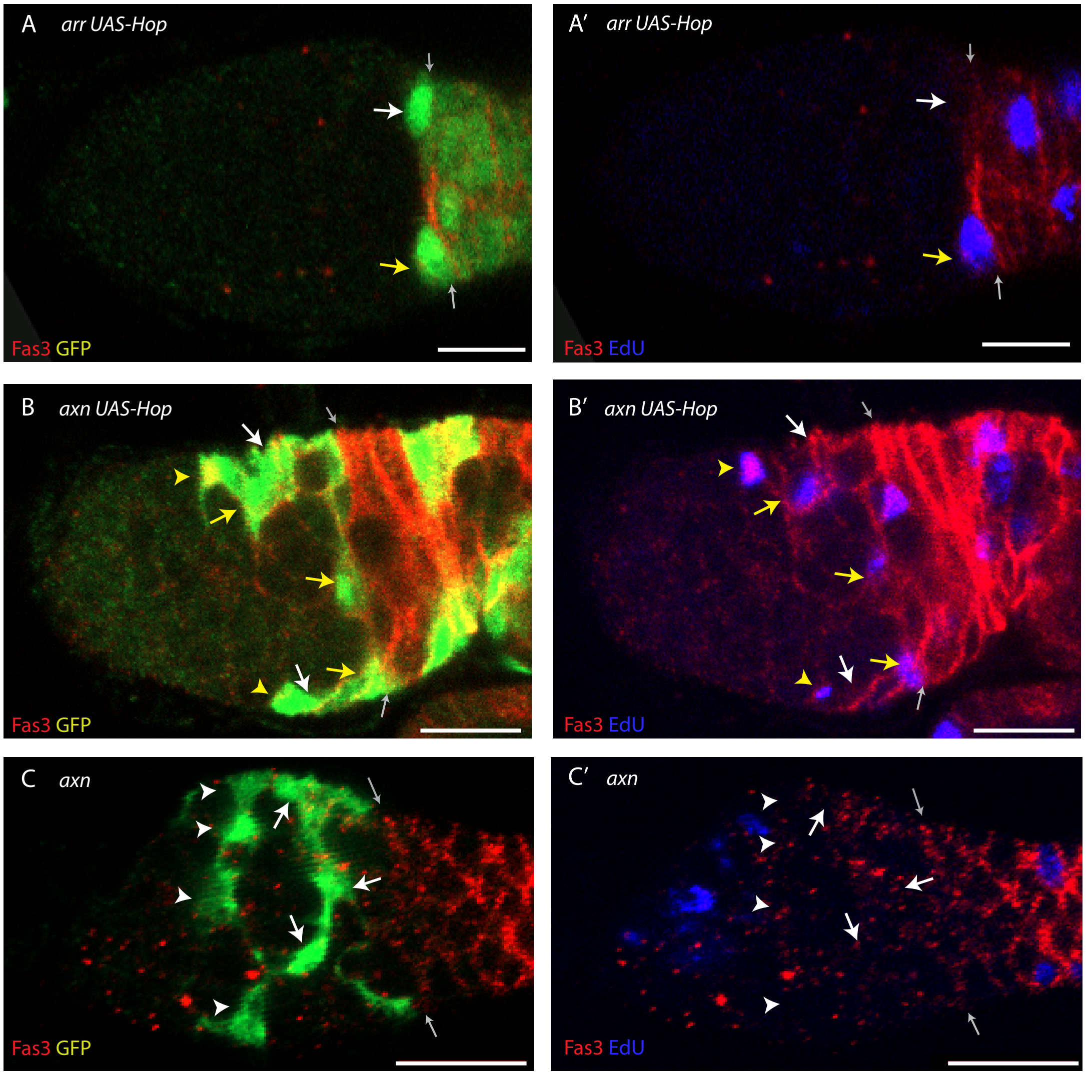
A arr UAS-Hop
Fas3 GFP
A′ arr UAS-Hop
Fas3 EdU
B axn UAS-Hop
Fas3 GFP
B′ axn UAS-Hop
Fas3 EdU
C axn
Fas3 GFP
C′ axn
Fas3 EdU

Figure S3

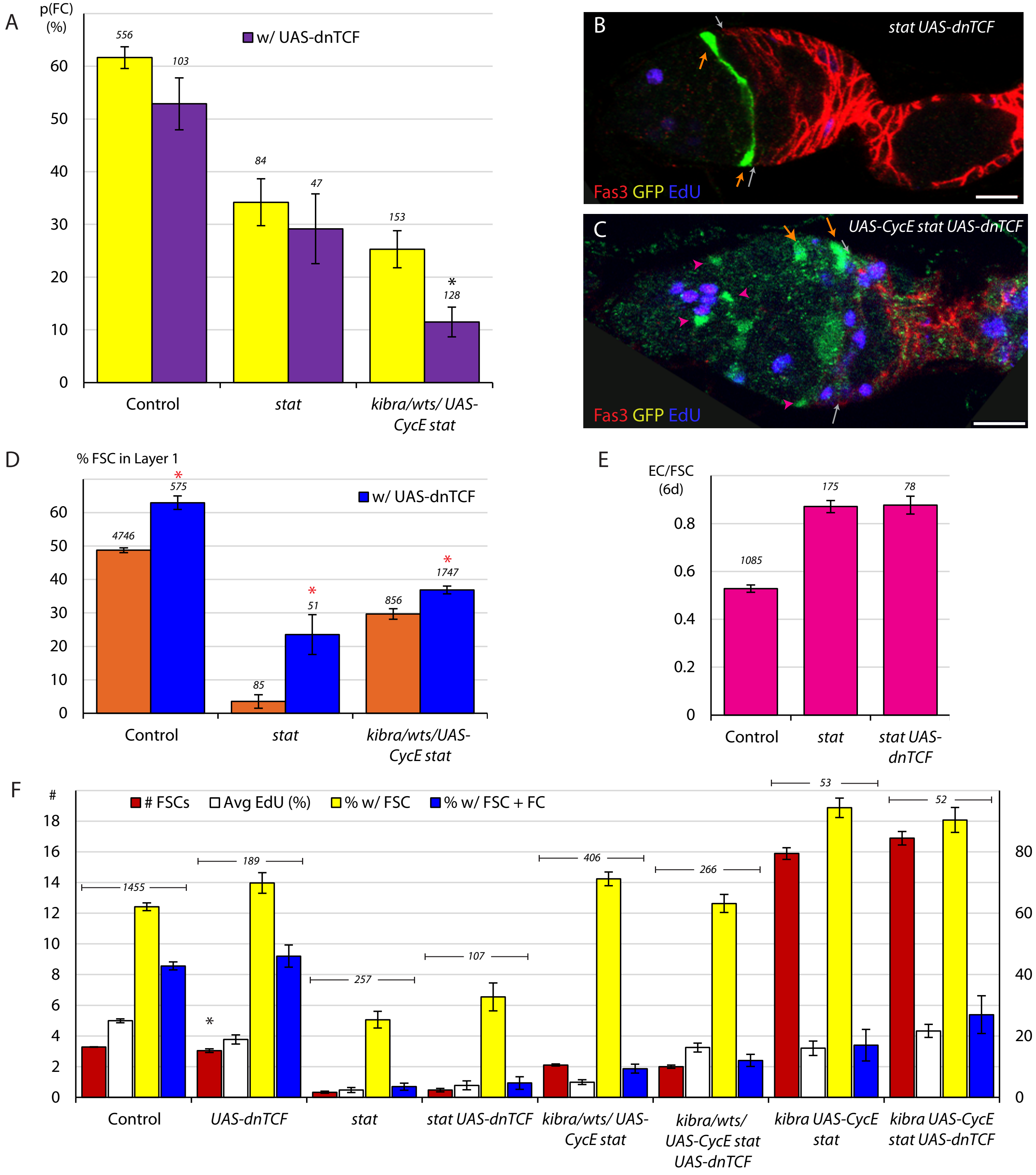
A
p(FC) (%)
w/ UAS-dnTCF
556
103
84
47
153
128
*
60
50
40
30
20
10
0
Control
stat
kibra/wts/ UAS-CycE stat
B
stat UAS-dnTCF
Fas3 GFP EdU
C
UAS-CycE stat UAS-dnTCF
Fas3 GFP EdU
D
% FSC in Layer 1
w/ UAS-dnTCF
4746
575
85
51
856
1747
*
Control
stat
kibra/wts/UAS-CycE stat
E
EC/FSC (6d)
1085
175
78
0.8
0.6
0.4
0.2
0
Control
stat
stat UAS-dnTCF
F
#
### FSCs
Avg EdU (%)
% w/ FSC
% w/ FSC + FC
1455
189
257
107
406
266
53
52
18
16
14
12
10
8
6
4
2
0
80
60
40
20
Control
UAS-dnTCF
stat
stat UAS-dnTCF
kibra/wts/ UAS-CycE stat
kibra/wts/ UAS-CycE stat UAS-dnTCF
kibra UAS-CycE stat
kibra UAS-CycE stat UAS-dnTCF

Figure S4
